# Lifestyle intervention is associated with attenuation of ER stress/inflammation and enhancement of naive immune cell identity in older adults with metabolic disease

**DOI:** 10.64898/2026.05.08.723786

**Authors:** Rushil Yerrabelli, Asa Thibodeau, Djamel Nehar-Belaid, Radu Marches, Sathyabaarathi Ravichandran, Yoann Barnouin, Ranjan Sen, Silke Paust, Michael L. Stitzel, Luigi Ferrucci, A. Phillip West, Jacques Banchereau, Dennis T. Villareal, George A. Kuchel, Duygu Ucar

**Affiliations:** The Jackson Laboratory for Genomic Medicine, Farmington, CT 06032, USA; Division of Endocrinology, Diabetes and Metabolism, Baylor College of Medicine, Houston, TX 77030, USA; Center for Translational Research on Inflammatory Diseases, Michael E DeBakey VA Medical Center, Houston, TX 77030, USA; Laboratory of Molecular Biology and Immunology, National Institute on Aging, Baltimore, MD 21224, USA; Intramural Research Program, National Institute on Aging, NIH, Baltimore, MD, 21224, USA; Department of Immunology, Tufts University School of Medicine, Boston, MA, 02111, USA; Mammalian Genetics, The Jackson Laboratory, Bar Harbor, ME, 04609, USA; Immunoledge LLC, Montclair, NJ 07042, USA; UConn Center on Aging, UConn Health, Farmington, CT 06030, USA; Institute for Systems Genomics, University of Connecticut Health Center, Farmington, CT 06030, USA; Department of Genetics and Genome Sciences, University of Connecticut Health Center, Farmington, CT 06030, USA

## Abstract

Chronic inflammation and cellular stress are hallmarks of aging, obesity, and type 2 diabetes (T2D), but whether these programs can be modulated by lifestyle intervention in late life, particularly in the presence of established metabolic disease, remain unknown. We profiled circulating immune cells from older adults with obesity and T2D (ages 66–83 years; *n* = 9) before and after a 6-month lifestyle intervention combining caloric restriction with exercise training. Participants showed substantial weight loss (∼7%) alongside improvements in glycemic control, insulin sensitivity, and physical performance. Longitudinal single-cell transcriptomic and epigenomic profiling identified two major changes. First, intervention was associated with downregulation of inflammatory and endoplasmic reticulum (ER) stress transcriptional programs, with the most pronounced effects observed in CD14^+^ monocytes. *DDIT3* (CHOP) was transcriptionally and epigenetically downregulated and its inferred regulatory network encompassed multiple inflammatory mediators. Second, naive CD4^+^ T, naive T_reg_, and naive B cells exhibited an upregulation of naive cell identity genes, with naiveness scores increasing after intervention, which declines with age in an independent healthy adult cohort. Together, these findings suggest that lifestyle intervention is associated with coordinated remodeling of both innate and adaptive immune compartments in older adults, revealing substantial plasticity of the aging immune system especially targeting ER stress, inflammation, and naive lymphocyte identity programs.

## Introduction

Aging is accompanied by a progressive remodeling of the immune system, characterized by chronic low-grade inflammation, accumulation of dysfunctional immune cells, and impaired immune responses^1–9^. These changes contribute to increased susceptibility to infection, impaired responses to vaccination, and elevated risk of chronic diseases in older adults^10–16^. Obesity and type 2 diabetes (T2D), which disproportionally affect older populations (40% of adults ≥60 years have obesity^17^ and 30% of adults ≥65 years have T2D^18^) further exacerbate immune aging by imposing additional inflammatory burden and metabolic stress^19–21^. Despite growing recognition of the immune consequences of aging, obesity, and T2D, whether these immune changes are reversible remains unknown^22^. This knowledge gap persists in part because older adults with established metabolic disease are often frail and therefore excluded from intervention trials^23,24^. The co-occurrence of obesity and age-related muscle loss, termed sarcopenic obesity, is particularly prevalent in this population and is associated with compounding risks of frailty, cardiometabolic dysfunction, and physical disability^25^.

Caloric restriction (CR) combined with aerobic and resistance exercise has been shown to improve insulin sensitivity, glycemic control, and physical performance in frail and pre-frail obese older adults^26–29^. Mechanistically, CR reduces nutrient-driven mTOR and inflammatory signaling, promotes autophagy, and shifts metabolism toward fatty acid oxidation and ketone body production, collectively dampening chronic inflammation^30–32^. In humans, the landmark CALERIE trial demonstrated that sustained CR in younger, metabolically healthy adults improved thymic function and induced anti-inflammatory transcriptional programs in adipose tissue^33^. Whether transcriptional remodeling extends to circulating immune cells, and whether similar molecular changes are achievable in older adults with established metabolic disease, are open questions.

To address these questions, we profiled peripheral blood mononuclear cells (PBMCs) from a cohort of older adults with obesity and T2D before and after a 6-month lifestyle intervention. This intervention, conducted as part of the Lifestyle Intervention for Seniors with Diabetes (LISD) trial, incorporated CR and exercise training, which resulted in substantial weight loss and improved metabolic and physical function^28^. Using longitudinal scRNA-seq and snATAC-seq, we show that lifestyle intervention is associated with coordinated attenuation of inflammatory and ER stress programs across circulating immune lineages, alongside an upregulation of naive lymphocyte identity programs, suggesting that the aging immune system retains plasticity even in the presence of advanced metabolic disease.

## RESULTS

### Lifestyle Intervention for Seniors with Diabetes (LISD) trial

As part of the LISD trial (ClinicalTrials.gov NCT02348801), community-dwelling older adults (65+ years) with obesity and T2D were enrolled in an intensive lifestyle intervention study consisting of CR (500–750 kcal/day deficit) combined with exercise^28^. Participants met as a group with dieticians, kept food diaries, and adjusted their dietary intake to match their prescribed balanced diets. They also participated in three supervised 90-minute exercise sessions per week including aerobic, resistance, and balance exercises. To investigate whether the metabolic and physical performance benefits observed in this cohort were accompanied by immune remodeling, we profiled PBMCs from nine participants (ages 66–83 years) collected before (baseline) and after the 6-month intervention (Fig. 1a). Frailty Index (FI)^34^ was calculated at the beginning of the intervention; seven of nine participants were frail (FI = 0.220-0.314), whereas two were prefrail (FI= 0.186-0.207) based on frailty level categories^35^. Further details of the study design and intervention are described in the parent study^28^.

**Fig. 1.**
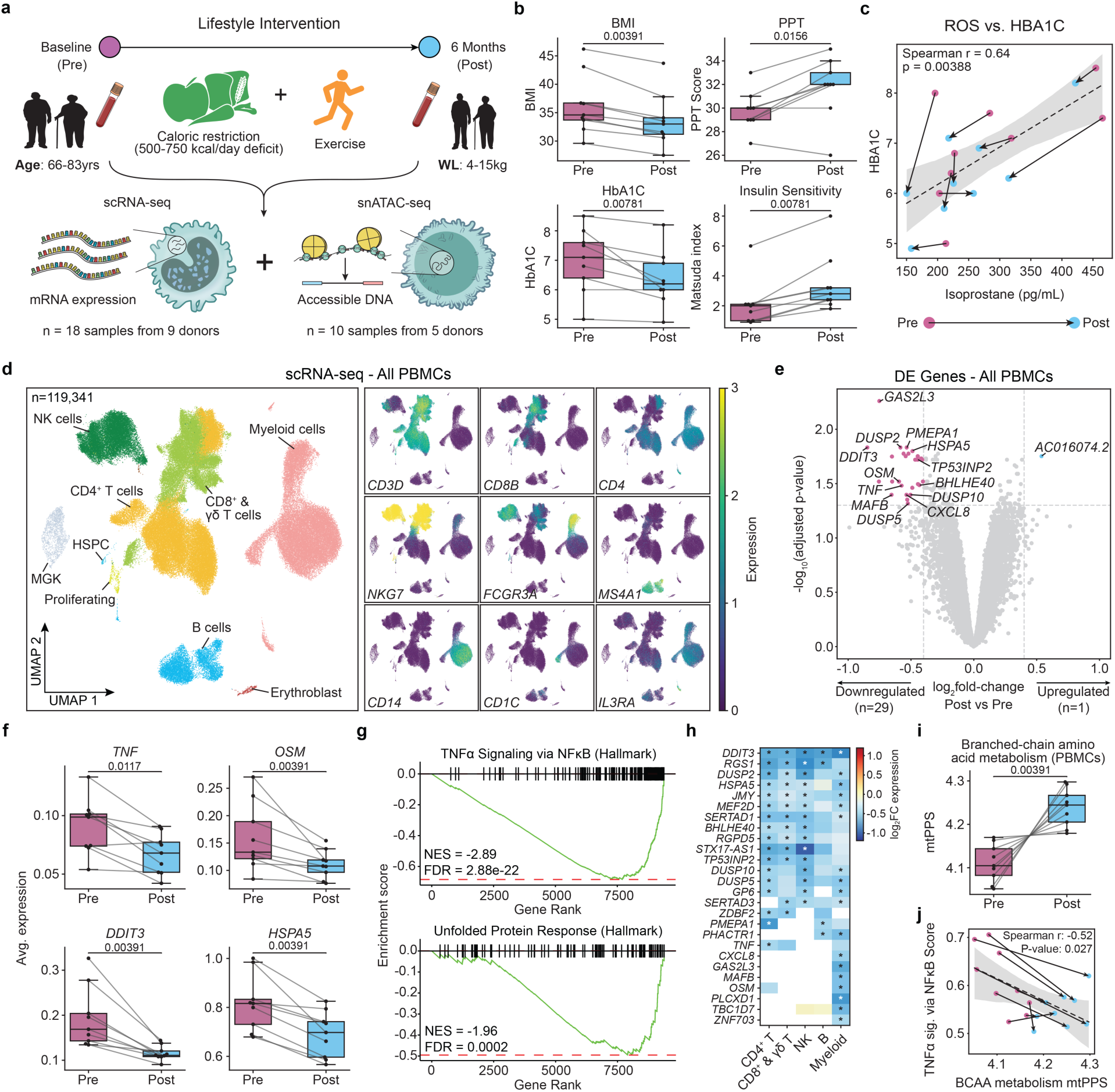
Lifestyle intervention is associated with downregulation of inflammatory and ER stress programs in circulating immune cells. **a**, Schematic of the study design. PBMCs were collected from nine older adults with obesity and type 2 diabetes (ages 66–83 years) before (Pre) and after (Post) a 6-month lifestyle intervention and profiled by scRNA-seq (n = 18 samples from 9 donors) and snATAC-seq (n = 10 samples from 5 donors). **b**, Paired comparisons of BMI, Physical Performance Test (PPT) score, HbA1c, and Matsuda index before and after intervention. **c**, Correlation between serum isoprostane levels and HbA1c across donor timepoints (pink, Pre; blue, Post). **d**, UMAP of 119,341 PBMCs colored by annotated cell population (left) and feature plots of canonical marker genes (right). **e**, Volcano plot of differentially expressed genes in total PBMCs (Post versus Pre). **f**, Average expression of selected differentially expressed genes before and after intervention. **g**, GSEA plots for Hallmark ’TNFα Signaling via NF-κB’ and ’Unfolded Protein Response’ gene sets in total PBMCs. Normalized enrichment score (NES) and FDR from GSEA are shown. **h**, Heatmap of log_2_ fold changes (Post versus Pre) for downregulated differentially expressed genes across major PBMC lineages. Asterisks indicate FDR < 5% within the corresponding lineage. **i**, BCAA metabolism mitochondrial pathway prioritization score (mtPPS) in total PBMCs before and after intervention. **j**, Correlation between BCAA metabolism mtPPS and ’TNFα Signaling via NF-κB’ leading-edge gene expression score across donor timepoints. Correlations and p values shown in c and j were calculated using Spearman’s correlation coefficient. P values in panels b and f and i were calculated using the Wilcoxon signed-rank test.

Clinical assessments confirmed metabolic and functional improvements following intervention (Fig. 1b, Table 1, Extended Data Fig. 1a), with detailed baseline characteristics and clinical responses provided in Supplementary Table 1). Participants exhibited significant reduction in body mass index (BMI; ∼7.2% decrease; p < 0.01), improved physical performance (Physical Performance Test (PPT); ∼6.6% increase; p < 0.05), improved insulin sensitivity (Matsuda Index; ∼1.3 unit increase; p < 0.01), and improved glycemic control (HbA1C; ∼0.62% decrease; p < 0.01) (Fig. 1b). We assessed oxidative stress by measuring serum F2-isoprostanes from serum, a gold standard biomarker of free radical–driven lipid peroxidation^36^. Isoprostane levels declined following intervention (p = 0.055), and this decline was significantly correlated with changes in HbA1c levels (Spearman r = 0.64, p < 0.01; Fig. 1c, Extended Data Fig. 1a). This is consistent with prior evidence that oxidative stress, as reflected by F2-isoprostane levels, tracks with glycemic burden in T2D^37^.

**Table 1.** Baseline Characteristics and Clinical Responses of Participants Included in Immune Profiling Analysis (n = 9)

| Variable | Value |
| --- | --- |
| Age (yr.) | 72.9 ± 1.7 |
| Sex, (male/female), n | 6/3 |
| Frailty index | 0.24 ± 0.01* |
| Body weight (kg) |  |
| Baseline | 104.9 ± 4.3 |
| Change at 6 months | -7.6 ± 1.1† |
| Body mass index (kg/m<sup>2</sup>) |  |
| Baseline | 36.3 ± 1.8 |
| Change at 6 months | -2.2 ± 0.4† |
| Insulin sensitivity (Matsuda index) |  |
| Baseline | 2.1 ± 0.6 |
| Change at 6 months | 1.3 ± 0.3† |
| HbA1c (%) |  |
| Baseline | 7.0 ± 0.3 |
| Change at 6 months | -0.6 ± 0.1† |
| Physical Performance Test (score) ‡ |  |
| Baseline | 29.8 ± 0.5 |
| Change at 6 months | 2.0 ± 0.5† |

Values are mean ± SE unless otherwise indicated.
\*Frailty index calculated according to the Rockwood deficit accumulation approach; values ≥0.21 indicate frailty<sup>34</sup>.
†Change from baseline to 6 months was assessed using two-sided Wilcoxon signed-rank tests: body weight (P = 0.0077), body mass index (P = 0.0078), insulin sensitivity index (P = 0.0078), HbA1c (P = 0.0078), and Physical Performance Test (P = 0.0156).
‡Physical Performance Test score range 0–36; higher scores indicate better physical performance<sup>89</sup>.
Clinical responses in this subgroup were consistent with those reported in the parent trial and are shown here for context<sup>28</sup>.

### Intervention is associated with downregulation of inflammation and ER stress genes in PBMCs

Transcriptional and epigenetic changes associated with intervention were mapped using single-cell RNA sequencing (scRNA-seq; n=18 samples from 9 donors) and single-nucleus ATAC-sequencing (snATAC-seq; n=10 samples from 5 donors). After quality control, doublet removal, and batch correction, scRNA-seq and snATAC-seq yielded 119,341 and 39,777 high-quality cells and nuclei respectively for downstream analyses (Fig. 1d, Extended Data Fig. 1b and c). Expression of canonical marker genes were used to annotate clusters into eight major populations: CD4^+^ T cells (*CD3D*, *CD4*), CD8^+^ T and γδ T cells (*CD3D*, *CD8B*), NK cells (*NKG7*), B cells (*MS4A1*), myeloid cells (*CD14, CD1C*), proliferating cells (*MKI67*), megakaryocytes (MGK; *PPBP*), hematopoietic stem and progenitor cells (HSPCs) (*CD34*), and erythroblasts (*ALAS2*) (Fig. 1d; Extended Data Fig. 1d). Annotations from scRNA-seq were transferred onto the snATAC-seq data using label transfer and used to annotate snATAC-seq clusters alongside gene activity scores (Methods; Extended Data Fig. 2a and b).

Although the intervention did not result in significant changes in the cellular composition of these major populations (Extended Data Fig. 2c; Supplementary Table 2), differential expression analysis of total PBMCs identified one upregulated and 29 downregulated genes following the intervention (FDR < 5%; Fig. 1e; Supplementary Table 3). Downregulated genes included inflammatory *TNF* and *OSM,* and unfolded protein and endoplasmic reticulum (ER) stress response-associated *HSPA5* (BiP) and *DDIT3* (CHOP) genes (Fig. 1f). Gene set enrichment analysis (GSEA)^38,39^ showed that ‘TNFα signaling via NF-κB’ (normalized enrichment score (NES) = -2.89; FDR < 1%) and ‘Unfolded protein response’ (NES = -1.96; FDR < 1%) Hallmark gene sets^40^ were downregulated after intervention (Fig. 1g, Extended Data Fig. 2d; Supplementary Table 3). To determine whether these transcriptional changes were shared or lineage-specific, we repeated the differential expression analysis across major PBMC lineages. Notably, *DDIT3* was downregulated across all lineages (Fig. 1h), whereas pro-inflammatory *CXCL8* and *OSM* were uniquely downregulated in myeloid cells, and *TNF* was uniquely downregulated in CD4^+^ T cells.

Together, lifestyle intervention was associated with significant downregulation of ER stress and inflammation-associated genes at the PBMC level.

### Intervention is associated with increased branched-chain amino acid metabolism programs

To investigate the effects of intervention on mitochondrial metabolic pathways, we calculated mitochondrial pathway prioritization scores (mitoPPS)^41^ from total PBMC expression. The top-ranked mitochondrial pathway was ‘Branched-chain amino acid (BCAA) metabolism’ (p < 0.01) (Extended Data Fig. 3a), whose dysregulation has been associated with obesity and insulin resistance^42^. Consistently, expression of *BCAT2* and *BCKDHB*, a subunit of the BCAA catabolic enzyme complex^43^, was increased after intervention (p < 0.05) (Extended Data Fig. 3b and c). Consistently, BCAA metabolism was correlated with insulin sensitivity and negatively correlated with the average expression of ‘TNFα signaling via NF-κB’ leading edge genes (Spearman r = 0.62 and -0.52 respectively; p < 0.05) (Fig. 1j, Extended Data Fig. 3d).

Repeating this analysis across PBMC lineages showed that most subsets exhibited significantly increased BCAA metabolism scores (p < 0.05), with B cells showing a similar trend (p = 0.0547) (Extended Data Fig. 3e). Further investigation of lineage subsets revealed that the most pronounced increases were observed in CD14^+^ monocyte and GZMK^+^ GZMH^+^ CD8⁺ T cell subsets (detailed in subsequent sections) (p_adj_ < 0.05), with similar but less pronounced effects across other subsets (Extended Data Fig. 3f and g). BCAA metabolism scores were negatively correlated with ’TNFα signaling via NF-κB’ leading-edge gene expression for both of these subsets (Spearman r = -0.54 and -0.55 respectively; p < 0.05) and positively correlated with insulin sensitivity (Spearman r = 0.59 and 0.56 respectively, p< 0.05). While elevated circulating BCAAs have been associated with pro-inflammatory myeloid states in obesity^44^, the transcriptional upregulation of BCAA metabolism genes observed here, alongside reduced inflammatory programs, may reflect enhanced intracellular BCAA utilization as a feature of improved immune metabolic fitness.

### Intervention is associated with reduced inflammation and enhanced naive cell identity in CD4+ T cells

CD4^+^ T cells clustered into four major subsets: naive CD4^+^ T (*CCR7*; n = 11,620), memory CD4^+^ T (*S100A4*; n = 16,880), CD4^+^ T_EMRA_ (*NKG7*; n = 6,738), and T_reg_ cells (*FOXP3*; n = 2,470) (Fig. 2a). Although the intervention did not result in significant changes in the cellular frequencies Extended Data Fig. 4a; Supplementary Table 2), differential expression analysis identified 74 upregulated and 196 downregulated genes in naive CD4^+^ T cells; 7 upregulated and 34 downregulated genes in T_reg_ cells; 31 upregulated and 133 downregulated genes in memory CD4^+^ T cells; and 10 upregulated and 41 downregulated genes in CD4^+^ T_EMRA_ cells (FDR < 5%; Extended Data Figure 4b; Supplementary Table 3). GSEA showed significant downregulation of ‘TNFα signaling via NFKB’ pathway genes including *TNF*, *NFKB1*, and *RELB*. Downregulation of *TNF* was driven by CD4^+^ T_EMRA_ cells, while *NFKB1* and *RELB* were broadly downregulated in naive and memory CD4^+^ T cells. ER stress-related genes (e.g., *HSPA5* and *DDIT3*) and ‘MAPK signaling’ genes (e.g. *DUSP2*) were broadly downregulated across CD4^+^ T cell subsets (Fig. 2b, Extended Data Fig. 4c). Genes associated with ’mTORC1 signaling’ were downregulated across CD4+ T cell subsets (Fig. 2b), including *CXCR4*, whose expression in naive T cells is regulated by mTOR signaling and linked to naive T cell homeostasis and promoting naive and stem-like T cell states in humans^45,46^.

**Fig. 2.**
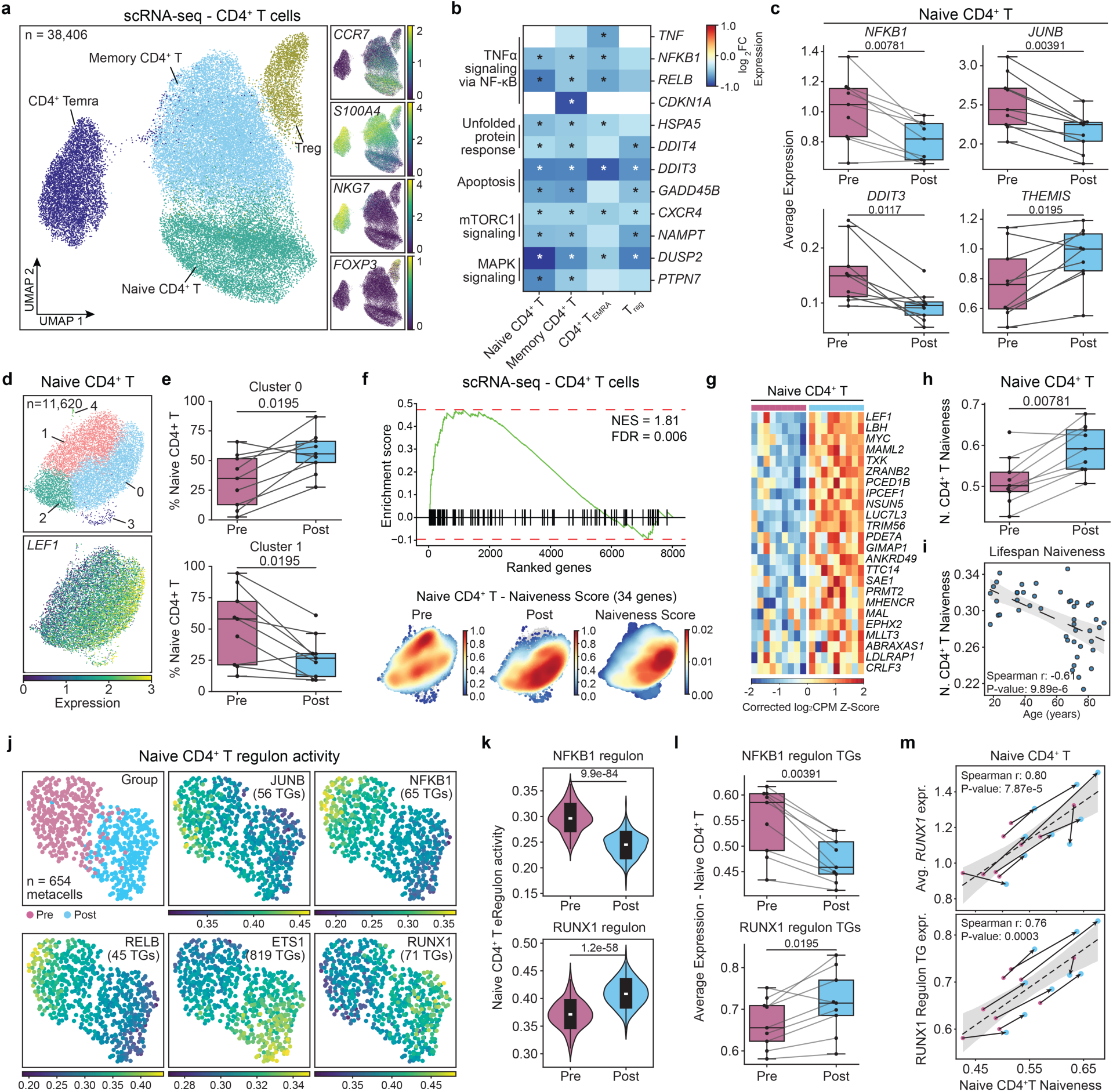
Lifestyle intervention is associated with downregulation of inflammatory programs and upregulation of naive cell identity in CD4⁺ T cells. **a**, UMAP of 38,406 CD4⁺ T cells colored by annotated subset (left) and feature plots of canonical marker genes (right). **b**, Heatmap of log_2_ fold changes (Post versus Pre) for selected differentially expressed genes across CD4⁺ T cell subsets that were among significantly downregulated gene sets from GSEA. Asterisks indicate FDR < 5% based on differential expression analysis within the subset. **c**, Average expression of selected genes in naive CD4⁺ T cells before and after intervention. **d**, UMAP of naive CD4⁺ T cells (n = 11,620) colored by cluster (top) and LEF1 expression (bottom). **e**, Frequencies of Cluster 0 and Cluster 1 as a percentage of naive CD4⁺ T cells before and after intervention. **f**, GSEA plot for the naive CD4⁺ T cell identity gene set (top) and density plots of pre and post intervention cells, and naiveness scores in naive CD4⁺ T cells (bottom). Normalized enrichment score (NES) and FDR from GSEA is shown. **g**, Heatmap of corrected log_2_ CPM z-scores for select naiveness leading-edge genes across donors before and after intervention. **h**, Naiveness scores in naive CD4⁺ T cells before and after intervention. **i**, Correlation between naive CD4⁺ T cell naiveness scores and age in healthy adults (>18 years, n=45). **j**, UMAP of SCENIC+ metacells (n = 654) colored by group and regulon activity for JUNB, NFKB1, RELB, ETS1, and RUNX1. **k**, Regulon activity for NFKB1 (top) and RUNX1 (bottom) regulons in naive CD4⁺ T cells before and after intervention. **l**, Average expression of NFKB1 (top) and RUNX1 (bottom) regulon target genes in naive CD4⁺ T cells before and after intervention. **m**, Correlations between naiveness scores and RUNX1 expression (top) and RUNX1 regulon target gene expression (bottom) in naive CD4⁺ T cells. Correlations and p values shown in i and m were calculated using Spearman’s correlation coefficient. P values in c, e, h, and l were calculated using the Wilcoxon signed-rank test. P values in k were calculated using the Mann–Whitney U test.

Consistent with this, *THEMIS*, a T cell development gene^47,48^, was among the top upregulated genes in naive CD4⁺ T cells (Fig. 2c, Extended Data Fig. 4d), prompting further characterization of this subset. Clustering identified five naive CD4⁺ T cell clusters (Fig. 2d; C0-C4), of which the frequency of C0, marked by high expression of *LEF1*, which highly expressed in human naive T cells and is downregulated upon antigen encounter^49^, was significantly increased following intervention (p < 0.05) (Fig. 2e; Extended Data Fig. 4e). To systematically assess whether intervention enhanced naive T cell identity programs more broadly, we performed GSEA using a ’Naiveness’ gene set (n=95 genes, Methods; Supplementary Table 5) defined from genes highly expressed in naive CD4⁺ T cells compared to other CD4⁺ T cell subsets in an independent cohort of 95 healthy individuals^3^. Genes upregulated after intervention were significantly enriched for this gene set (NES = 1.81, FDR < 5%) (Fig. 2f; Supplementary Table 4), with leading-edge genes including *LEF1* and *GIMAP1*, a regulator of T cell survival^50^ (Fig. 2g). ’Naiveness score’, the mean expression of these leading-edge genes, was significantly higher in naive CD4⁺ T cells following intervention (p < 0.05) (Fig. 2h). Strikingly, the same score declined with age in healthy adults (>18 years, n=45) (Spearman *r* = −0.61, *p* < 0.001), suggesting that the intervention partially reverses age-associated transcriptional changes in naive CD4⁺ T cells (Fig. 2i).

Integration of scRNA-seq and snATAC-seq data from naive CD4⁺ T cells using SCENIC+^51^ identified 30 regulons (n=654 meta-cells; Supplementary Table 6; Methods) linking transcription factor (TF) motif accessibility to target gene expression. These regulons distinguished pre- and post-intervention cell states (Fig. 2j-l, Extended Data Fig. 4f-i). Regulatory networks for NFKB1, RELB, and BACH2 were more active pre-intervention, consistent with the broad downregulation of inflammatory transcription programs. In contrast, regulatory network activity for RUNX1 and ETS1 was increased post-intervention, consistent with transcriptional features associated with naive cell homeostasis^52–55^ (Fig 2j and k). Concordantly, cumulative average expression of NFKB1 target genes was reduced, while RUNX1 and ETS1 target gene expression was increased after intervention (p < 0.05) (Fig 2l, Extended Data Fig. g). Expression of *RUNX1* and *ETS1*, as well as their respective target genes, correlated with naiveness scores in naive CD4⁺ T cells (Spearman r = 0.80 and 0.76 for RUNX1 and RUNX1 targets; Spearman r = 0.52 and 0.90 for ETS1 and ETS1 targets; p < 0.001) (Fig, 2m, Extended Data Fig. 4j), further supporting a coordinated shift toward a naive cell identity restoration.

T_reg_ cells clustered into naive (n=1,010) and memory (n=1,460) subsets, with frequencies unchanged by intervention (Extended Data Fig. 5a and b). GSEA revealed positive enrichment of naive cell identity genes in naive Tregs following intervention (NES = 1.95, FDR < 5%), with leading edge genes including *LEF1* and *TCF7*, of which 19 genes overlapped with those identified in naive CD4⁺ T cells (Extended Data Fig. 5c-e; Supplementary Table 4). “Treg naiveness score” derived from these leading-edge genes was significantly increased after intervention (p < 0.01) and declined with age in healthy adults (>18 years, n=45) (Spearman *r* = -0.36, p < 0.05), paralleling the age-associated pattern observed in naive CD4⁺ T cells (Extended Data Fig. 5f and g).

Lifestyle intervention was associated with coordinated remodeling of naive CD4⁺ T cells and naive Tregs, characterized by downregulation of inflammatory and ER stress programs and upregulation of naive cell identity associated regulatory programs.

### Intervention is associated with downregulation of inflammation and ER-stress genes in memory CD4⁺ T cells

Memory CD4^+^ T cells clustered into eight subsets: central memory (T_CM_; *CCR7*; n = 3,586),Tfh (*CXCR5*; n = 3,923), Th1 (*IFNG-AS1*; n = 2,868), Th2 (*PTGDR2*; n = 345), Th17 (*RORC*; n = 2,697), Th22 (*CCR10*; n = 425), HLA-DR^+^ memory CD4^+^ T (*HLA-DRA*; n=393), and effector memory-like (EM-like; n = 2,287) subsets (Fig. 3a, Extended Data Fig. 6a). Although cell frequencies of these populations were unchanged by intervention (Extended Data Fig. 6b; Supplementary Table 2), differential expression analysis identified 136 downregulated and 42 upregulated genes in CD4^+^ T_CM_, 201 downregulated and 61 upregulated genes in Tfh, 109 downregulated and 22 upregulated genes in T_h_1, 18 downregulate and 2 upregulated genes in T_h_17, and 78 downregulated and 13 upregulated genes in CD4^+^ T_EM_-like subsets (Supplementary Table 3).

**Fig. 3.**
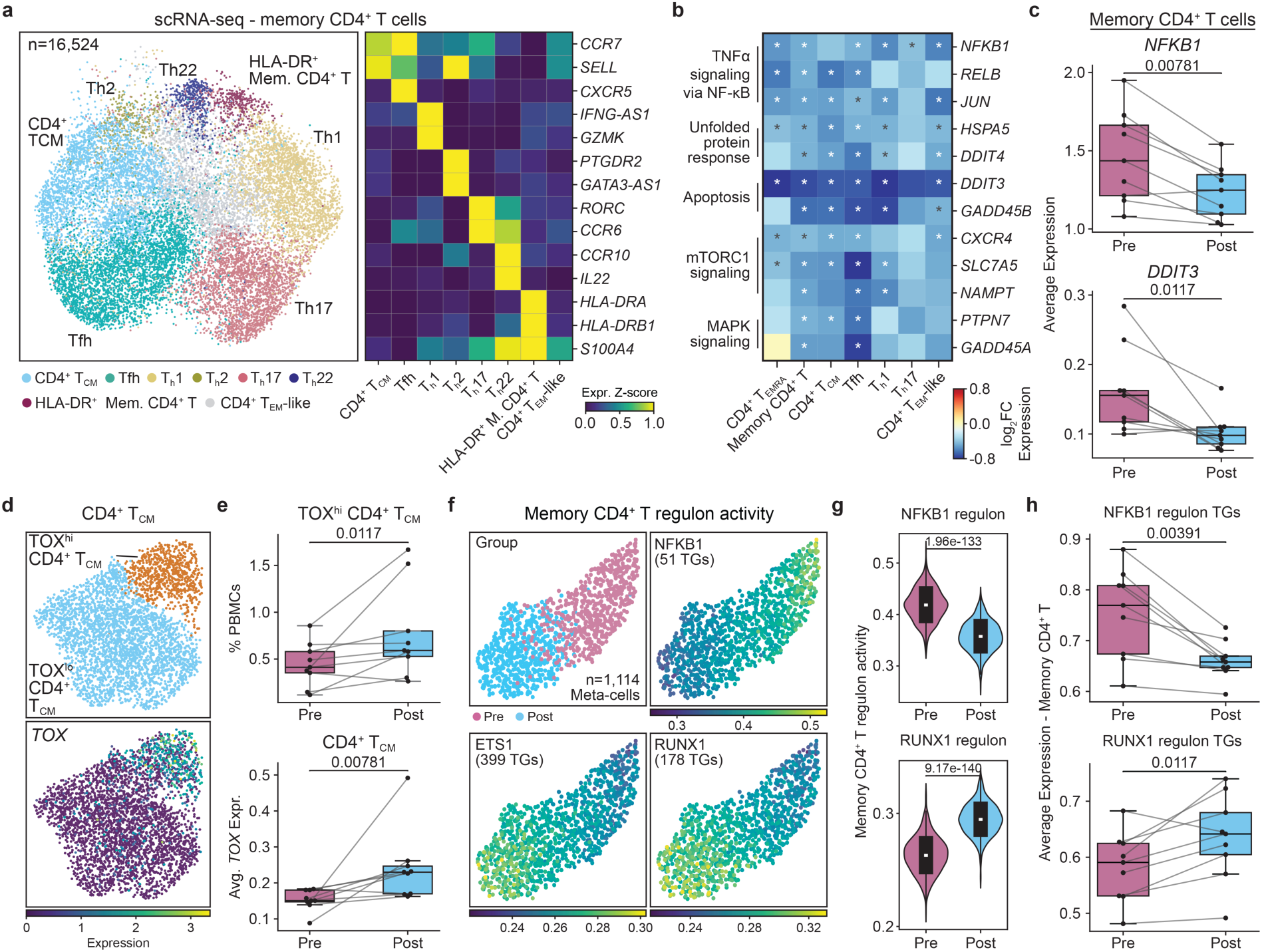
Lifestyle intervention is associated with downregulation of inflammatory programs and upregulation of TOX in memory CD4⁺ T cells. **a**, UMAP of 16,524 memory CD4⁺ T cells colored by annotated subset (left) and heatmap of marker gene expression z-scores used for annotation (right). **b**, Heatmap of log_2_ fold changes (Post versus Pre) for selected differentially expressed genes across memory CD4⁺ T cell subsets that were among significantly downregulated gene sets from GSEA. Asterisks indicate FDR < 5% based on differential expression analysis within the subset. **c**, Average expression of *NFKB1* and *DDIT3* in memory CD4⁺ T cells before and after intervention. **d**, UMAP of CD4⁺ TCM cells colored by TOX^hi^ and TOX^lo^ cluster assignments (top) and TOX expression (bottom). **e**, Frequency of TOX^hi^ CD4⁺ TCM cells as a percentage of total PBMCs (top) and average TOX expression in CD4⁺ TCM cells (bottom) before and after intervention. **f**, UMAP of SCENIC+ metacells (n = 1,114) colored by group and regulon activity for NFKB1, ETS1, and RUNX1. g, Regulon activity for NFKB1 (top) and RUNX1 (bottom) regulons in memory CD4⁺ T cells before and after intervention. **h**, Average expression of NFKB1 (top) and RUNX1 (bottom) regulon target genes in memory CD4⁺ T cells before and after intervention. P values in c, e, and h were calculated using the Wilcoxon signed-rank test. P values in g were calculated using the Mann–Whitney U test.

GSEA showed downregulation of ‘TNFα signaling *via* NF-κB’, ‘Unfolded protein response’, ‘Apoptosis’, ‘mTORC1 signaling’, and ‘MAPK signaling’ gene sets across memory CD4^+^ T cell subsets, including significant downregulation of *NFKB1*, *DDIT3*, *HSPA5*, and *CXCR4* (Fig. 3b and c, Extended Data Fig. 6c; Supplementary Table 4). In contrast, *TOX* was uniquely upregulated in CD4^+^ T_CM_ cells (Extended Data Fig. 6d). Further clustering of CD4^+^ T_CM_ cells identified a TOX^hi^ cluster, frequency of which increased after intervention (p < 0.01) (Fig. 3d and e). *TOX* has recently been associated with enhanced effector function in CD4⁺ T cells, distinct from its well-established and canonical role in CD8⁺ T cell exhaustion^56,57^. Exhaustion-related gene sets were not significantly enriched among upregulated genes (unadjusted p > 0.77) (Extended Data Fig. 6e), suggesting that *TOX* upregulation in this context may reflect enhanced effector function rather than exhaustion.

Integration of scRNA-seq and snATAC-seq data from memory CD4^+^ T cells using SCENIC+^51^ identified 58 regulons (n=1,114 meta-cells; Supplementary Table 6) that distinguished pre- and post-intervention cells (Fig. 3f). Regulatory networks for NFKB1 were more active pre-intervention, while RUNX1 and ETS1 regulatory activitiy was higher post-intervention (Fig. 3f-h, Extended Data Fig. f-i). Concordantly, cumulative average expression of NFKB1 target genes was reduced while RUNX1 and ETS1 target gene expression was increased after intervention (p < 0.05; Fig. 3h), paralleling the signatures observed in naive CD4^+^ T cells.

We examined chromatin accessibility at inferred TF binding sites using ChromVAR^58^ (Methods). Interestingly, accessibility of interferon regulatory factor (IRF) binding sites was among the top ranked TFs with respect to multiple statistical test p-values (Extended Data Fig. 7a; Supplementary Table 7; Methods). Among all IRF family members, only *IRF3* showed significant upregulation post-intervention (p < 0.05), an effect specific to Tfh cells (Extended Data Fig. b and c). *IRF3* expression was negatively correlated with *NFKB1* and positively correlated with *RUNX1* expression in memory CD4^+^ T cells (Extended Data Fig. d), consistent with transcriptional states characterized by lower inflammatory and higher naive identity-associated signatures.

Lifestyle intervention is associated with coordinated attenuation of inflammatory and ER stress transcriptional programs alongside increased expression of *IRF3* and naive identity-associated regulatory activity in memory CD4⁺ T cells.

### Intervention is associated with downregulation of inflammation and ER stress programs in CD8^+^ T and NK cells

CD8⁺ T and γδ T cells clustered into 8 subsets: naive CD8⁺ T cells (*CCR7* and *SELL*; n=1,322), GZMK⁺ CD8⁺ T cells (*GZMK*, n=4,079), GZMK⁺ GZMH⁺ CD8⁺ T cells (*GZMK* and *GZMH*; n=4,081), CD8⁺ T_EMRA_ cells (*GZMB*; n=6,713), KLRC2+ GZMK+ CD8+ T (KLRC2; n=666), MAIT (*ZBTB16*; n=571), γδ T cells (*TRDC* and *TRGC2*; n=2,779), and ISG^hi^ T cell (*ISG15*; n=167) populations (Fig. 4a). Although cell frequencies of these populations were unchanged by intervention (Extended Data Fig. 8a; Supplementary Table 2), differential expression analysis identified 1 downregulated gene in naive CD8^+^ T, 86 downregulated and 15 upregulated genes in GZMK^+^ CD8^+^ T cells, 46 downregulated and 11 upregulated genes in GZMK⁺ GZMH⁺ CD8⁺ T cells, 35 downregulated and 6 upregulated genes in CD8^+^ T_EMRA_ cells, 79 downregulated and 60 upregulated genes in γδ T cells, and 1 downregulated gene in MAIT cells (Supplementary Table 3).

**Fig. 4.**
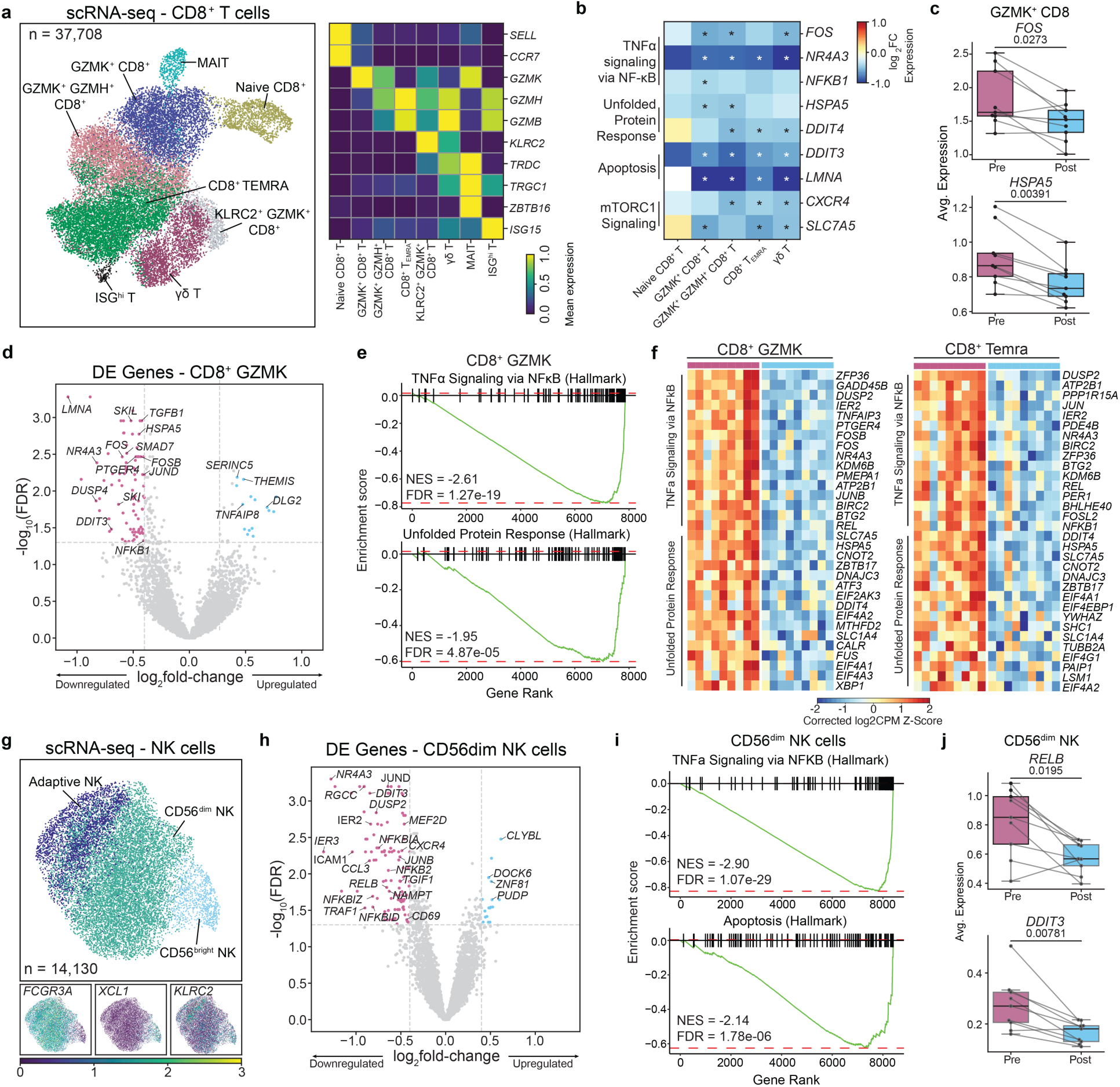
Lifestyle intervention is associated with downregulation of inflammatory and ER stress programs in CD8⁺ T and NK cells. **a**, UMAP of 37,708 CD8⁺ T cells colored by annotated subset (left) and heatmap of marker gene expression used for annotation (right). **b**, Heatmap of log_2_ fold changes (Post versus Pre) for selected differentially expressed genes across CD8⁺ T cell subsets that were among significantly downregulated gene sets from GSEA. Asterisks indicate FDR < 5% based on differential expression analysis within the subset. **c**, Average expression of *REL* and *HSPA5* in GZMK⁺ CD8⁺ T cells before and after intervention. **d**, Volcano plot of differentially expressed genes in GZMK⁺ CD8⁺ T cells (Post versus Pre). **e**, GSEA plots for Hallmark ’TNFα Signaling via NF-κB’ and ’Unfolded Protein Response’ gene sets in GZMK⁺ CD8⁺ T cells. Normalized enrichment score (NES) and FDR from GSEA are shown. **f**, Heatmaps of corrected log2 CPM z-scores for ’TNFα signaling via NF-κB’ and ’Unfolded protein response’ leading-edge genes in GZMK⁺ CD8⁺ T cells (left) and CD8⁺ T_EMRA_ cells (right) across donors before and after intervention. **g**, UMAP of 14,130 NK cells colored by annotated subset (top) and feature plots of canonical marker genes (bottom). **h**, Volcano plot of differentially expressed genes in CD56^dim^ NK cells (Post versus Pre). **i**, GSEA plots for Hallmark ’TNFα signaling via NF-κB’ and ’Apoptosis’ gene sets in CD56^dim^ NK cells. NES and FDR shown. **j**, Average expression of *RELB* and *DDIT3* in CD56^dim^ NK cells before and after intervention. P values in c and j were calculated using the Wilcoxon signed-rank test.

GSEA showed downregulation of ‘TNFα signaling via NFKB’, ‘Apoptosis’, ‘mTORC1 signaling’, and ‘Unfolded protein response’ gene sets in GZMK⁺ CD8⁺ T, GZMK⁺ GZMH⁺ CD8⁺ T, and CD8⁺ T_EMRA_ cells (p_adj_ < 0.05) (Fig. 4b, Extended Data Fig. 8b). GZMK⁺ CD8⁺ T cells exhibited the most prominent transcriptional changes among CD8^+^ T cells, including downregulation of *FOS*, *DDIT3*, and *HSPA5* (Fig. 4c,d). Downregulated genes were enriched for ‘TNFα signaling via NF-κB’ (NES = −2.61, FDR < 1%) and ‘Unfolded protein response’ (NES = −1.95, FDR < 1%) (Fig. 4e; Supplementary Table 4), with leading-edge genes including canonical inflammatory mediators *TNFAIP3*, *PTGER4*, *FOS*, *JUNB*, and NF-κB-related genes, alongside ER stress-associated genes *DDIT4*, *HSPA5*, *EIF2AK3*, and *XBP1* (Fig. 4f). CD8⁺ T_EMRA_ cells also showed concordant downregulation of ‘TNFα signaling via NF-κB’ (NES = -2.74, FDR < 1%) and ‘Unfolded protein response’ (NES = -1.70, FDR = < 5%) (Supplementary Table 4), with leading-edge genes including *JUN*, *REL*, *NFKB1*, *DDIT4*, and *HSPA5* (Fig. 4f).

NK cells clustered into CD56^bright^ NK (*NCAM1*; n=1,010), CD56^dim^ NK (*FCGR3A*; n=10,108), and adaptive NK (*KLRC2*; n=3,012) cell subsets (Fig. 4g), with cell frequencies unchanged by intervention (Extended Data Fig. 8c; Supplementary Table 2). Differential expression analysis identified 106 downregulated and 18 upregulated genes in CD56^dim^ NK cells, 63 downregulated and 3 upregulated genes in CD56^bright^ NK, and 3 downregulated genes in adaptive NK cells (Fig. 4h; Supplementary Table 3). GSEA showed downregulation of ‘TNFα signaling via NF-κB’ and ‘mTORC1 signaling’ gene sets across all NK cell subsets (FDR < 5%) (Fig. 4i, Extended Data Fig. 8d; Supplementary Table 4), including downregulation of *RELB* and *DDIT3* (p < 0.05) (Fig. 4j).

Together, lifestyle intervention was associated with downregulation of inflammatory and ER stress genes across CD8⁺ T and NK cell compartments.

#### Lifestyle intervention is associated with downregulation of inflammation and ER stress programs and upregulation of naive identity genes in B cells

B cells clustered into 5 subsets: naive B (*IGHD*; n = 3,860), transitional B (TrB; *MME*; n = 616), memory B (*CD27*; n = 2,219), age-associated B cells (ABCs, also referred to as atypical B cells; *ZEB2*; n = 540), and plasmablasts (*JCHAIN*; n = 67) (Fig. 5a). Although cell frequencies of these populations were unchanged by intervention (Extended Data Fig. 9a; Supplementary Table 2), differential expression analysis identified 17 downregulated and 2 upregulated genes in naive B cells, and 17 downregulated and 1 upregulated gene in memory B cells (Fig. 5b; Extended Data Fig. 9b; Supplementary Table 3). GSEA showed downregulation of ‘TNFα signaling via NFκB’, ‘Apoptosis’, ’Unfolded protein response’, and ‘MAPK signaling’ gene sets in naive and memory B subsets (Fig. 5c and d, Extended Data Fig. 9c; Supplementary Table 4), Downregulated genes included ER stress-associated genes *DDIT3*, *DDIT4*, and *HSPA5*, and inflammatory *NFKB1*, *REL*, *JUN*, and *FOS*.

**Fig. 5.**
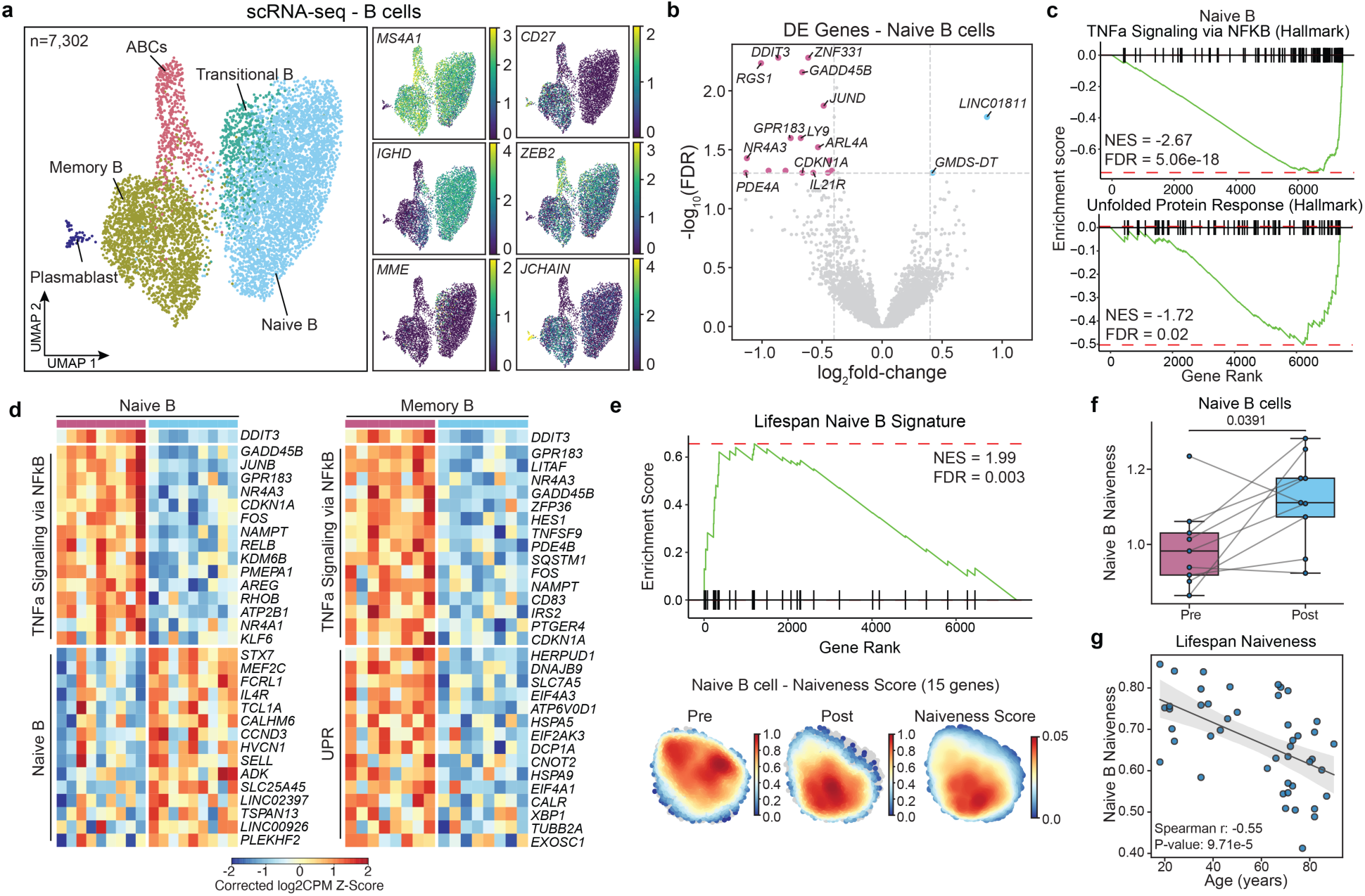
Lifestyle intervention is associated with downregulation of inflammatory programs and upregulation of naive cell identity in B cells. **a**, UMAP of 7,302 B cells colored by annotated subset (left) and feature plots of canonical marker genes (right). **b**, Volcano plot of differentially expressed genes in naive B cells (Post versus Pre). **c**, GSEA plots for Hallmark ’TNFα signaling via NF-κB’ and ’Unfolded protein response’ gene sets in naive B cells. Normalized enrichment score (NES) and FDR from GSEA are shown. **d**, Heatmaps of corrected log_2_ CPM z-scores for ’TNFα signaling via NF-κB’ leading-edge genes and naive B cell identity genes in naive B cells (left) and ’TNFα signaling via NF-κB’ and ’Unfolded protein response’ leading-edge genes in memory B cells (right) across donors before and after intervention. **e**, GSEA plot for the naive B cell identity gene set (top) and density plots of pre and post intervention cells, and naiveness scores in naive B cells (bottom). NES and FDR is shown. **f**, Naiveness scores in naive B cells before and after intervention. P value was calculated using the Wilcoxon signed-rank test. **g**, Correlation between naive B cell naiveness scores and age in an independent healthy adult cohort. Correlations and p values were calculated using Spearman’s correlation coefficient.

To assess whether naive B cells exhibit increased naive cell identity gene expression after intervention, we performed GSEA using a naive B cell signature (Supplementary Table 5) derived from healthy individuals^3^. Genes upregulated upon intervention in naive B cells were significantly enriched for this gene set, with leading-edge genes including *IL4R*, and *SELL*, essential genes for naive B cell identity and functions^59,60^. (NES = 1.99, FDR < 1%) (Fig. 5d and e; Supplementary Table 4). ‘B cell naiveness’ score derived from these leading-edge genes was significantly higher post-intervention (p < 0.05) (Fig. 5f) and declined with age in healthy adults (>18 years; n=45) (Spearman *r* = −0.55; *P* < 0.001) (Fig. 5g), paralleling the age-associated pattern observed in naive CD4⁺ T and T^reg^ cells. The increased expression of naive B cell identity-associated genes suggests that intervention may partially reverses age-associated transcriptional changes in naive B cells.

Lifestyle intervention was associated with downregulation of inflammatory and ER stress programs alongside upregulation of naive B cell identity genes.

#### Intervention is associated with downregulation of inflammation and ER stress programs in myeloid cells

Myeloid cells clustered into 5 subsets: CD14^+^ monocyte (*CD14*; n = 22,564), CD16^+^ monocyte (*FCGR3A*; n = 5,495), CD14^+^ CD16^+^ monocyte (*CD14* and *FCGR3A*; n = 1,511), DC (*CLEC10A* and *CLEC9A*; n=2,017), and pDC (*IL3RA* and *IRF7*; n=465) (Fig. 6a, SFig. 5a). Although cell frequencies of these populations were unchanged by intervention (Extended Data Fig. 10a; Supplementary Table 2), differential expression analysis identified 96 downregulated and 41 upregulated genes in CD14^+^ monocytes (Fig. 6b); 20 downregulated genes in CD16^+^ monocytes, 27 downregulated and 2 upregulated genes in CD14^+^ and CD16^+^ monocytes, and 8 downregulated and 2 upregulated genes in DCs (FDR < 5%; ; Supplementary Table 3).

**Fig. 6.**
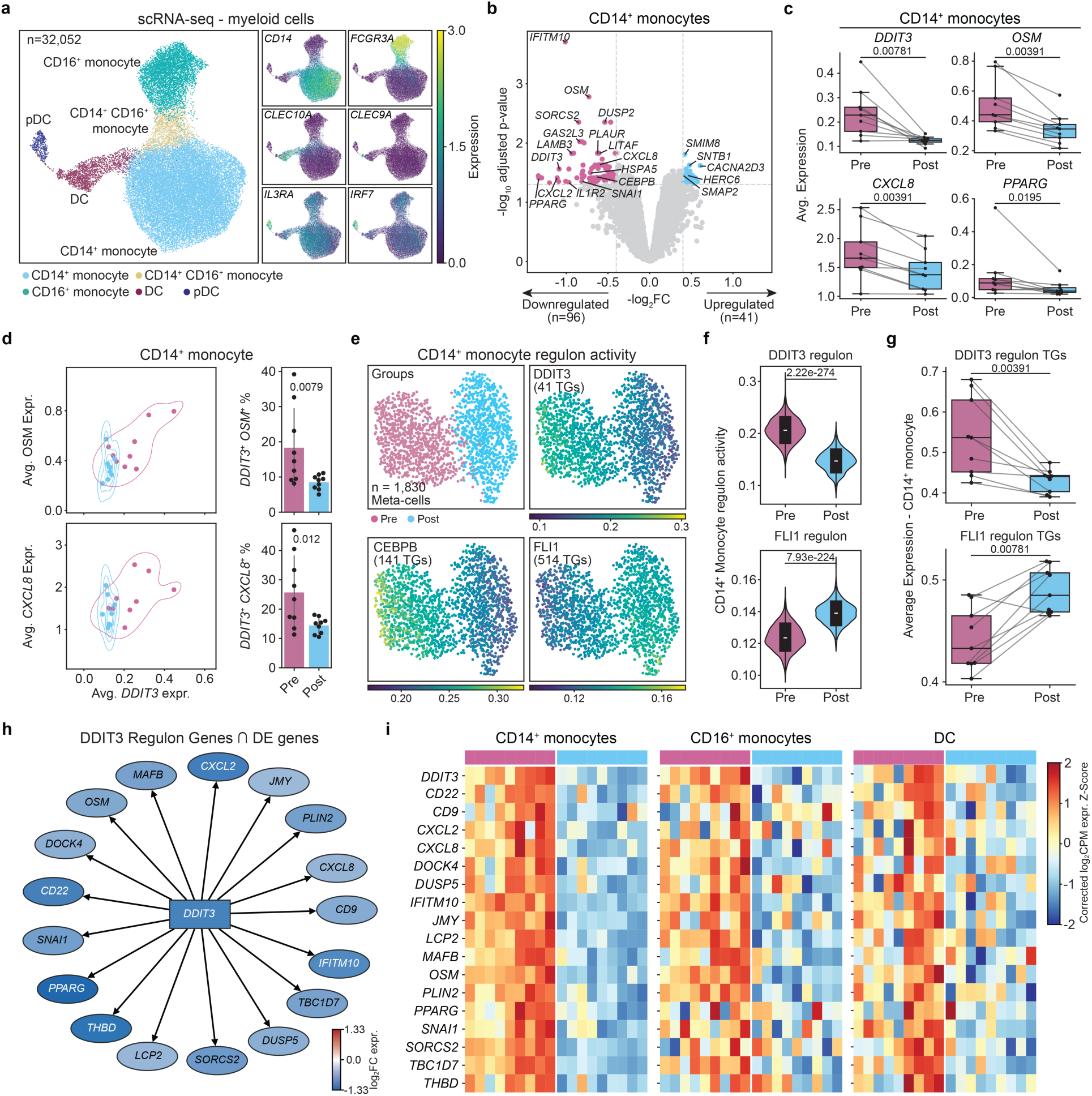
Lifestyle intervention is associated with downregulation of inflammatory and ER stress programs in myeloid cells, centered on DDIT3. **a**, UMAP of 32,052 myeloid cells colored by annotated subset (left) and feature plots of canonical marker genes (right). **b**, Volcano plot of differentially expressed genes in CD14⁺ monocytes (Post versus Pre). **c**, Average expression of *DDIT3*, *OSM*, *CXCL8*, and *PPARG* in CD14⁺ monocytes before and after intervention. **d**, Scatterplots of average *DDIT3* expression versus average *OSM* (top left) and *CXCL8* (bottom left) expression in CD14⁺ monocytes across donors (pink, Pre; blue, Post), and frequencies of DDIT3⁺OSM⁺ (top right) and DDIT3⁺CXCL8⁺ (bottom right) double-positive cells before and after intervention. Error bar shows represents the standard deviation of percentages across samples for each group. **e**, UMAP of SCENIC+ meta-cells (n = 1,830) colored by group and regulon activity for DDIT3, CEBPB, and FLI1. **f**, Regulon activity for DDIT3 (top) and FLI1 (bottom) regulons in CD14⁺ monocytes before and after intervention. P values in f were calculated using the Mann–Whitney U test. **g**, Average expression of DDIT3 (top) and FLI1 (bottom) regulon target genes in CD14⁺ monocytes before and after intervention. **h**, Network diagram of DDIT3 regulon target genes overlapping with differentially expressed genes in CD14⁺ monocytes, colored by log_2_ fold change. **i**, Heatmaps of corrected log_2_ CPM z-scores for DDIT3 regulon target gene overlap with CD14+ monocyte differentially expressed genes across donors in CD14⁺ monocytes, CD16⁺ monocytes, and DCs before and after intervention. P values in c, d, and g were calculated using the Wilcoxon signed-rank test.

Downregulated genes in CD14^+^ monocytes included *DDIT3*, the metabolic regulator *PPARG*, and inflammatory mediators *OSM* and *CXCL8* (Fig. 6b and c). Notably, *DDIT3* was co-expressed with *OSM* and *CXCL8* in within the same single CD14^+^ monocytes, and the number of double-positive cells (*DDIT3*^+^*OSM*^+^ and *DDIT3*^+^*CXCL8*^+^) was reduced after intervention (p < 0.05) (Fig. 6d), suggesting coordinated downregulation of ER stress and inflammatory programs at the single cell level.

GSEA showed downregulation of ‘TNFα signaling via NFκB’ gene set across all myeloid subsets (Extended Data Fig. 10b; Supplementary Table 4). This was alongside the downregulation of ‘mTORC1 signaling’, ‘oxidative phosphorylation’, ‘Apoptosis’, and ‘Unfolded protein response’ gene sets which were more prominently associated with CD14^+^ monocytes and DCs (Extended Data Fig. 10b; Supplementary Table 4). Significantly downregulated genes in these pathways included ER-stress associated *DDIT3* and *HSPA5* and inflammation associated *LITAF* and *CEBPB* (Extended Data Fig. 10c).

Integration of scRNA-seq and snATAC-seq data from CD14^+^ monocytes using SCENIC+^51^ identified 115 regulons (n=1,830 meta-cells; Supplementary Table 6) that distinguished pre- and post-intervention cells (Fig. 6e, Extended Data Fig. 10d). Regulatory networks anchored by DDIT3 and CEBPB were more active pre-intervention, whereas the regulatory activity for FLI1, an ETS family member involved in myeloid lineage regulation^61^, was increased post-intervention (Fig. 6e and f, Extended Data Fig. 10d-f). Concordantly, cumulative average expression of DDIT3 target genes was reduced while FLI1 target gene expression was increased after intervention (p < 0.05; Fig. 6g, Extended Data Fig. 10f). DDIT3 target genes overlapped with 17 significantly downregulated genes, including *OSM* and *CXCL8*. Similarly, these DDIT3 target genes were downregulated with intervention in in CD16⁺ monocytes and DCs (Fig. 6h and i), further emphasizing the association between DDIT3 and inflammatory gene expression, alongside its downregulation in response to the intervention.

Myeloid cells exhibited the most pronounced downregulation of ER stress and inflammatory genes following intervention, with inactivation of *DDIT3* observed both transcriptionally and epigenetically, emerging as a key molecule linking these two processes.

## DISCUSSION

Older adults with obesity and T2D represent a high-risk population with disproportionate burdens of cardiovascular disease, infection susceptibility, and functional impairment^21^. Using longitudinal single-cell transcriptomic and epigenomic profiling, we show that a 6-month lifestyle intervention combining CR and exercise is associated with widespread remodeling of both innate and adaptive circulating immune cells in this population. The most consistent finding was a pan-lineage downregulation of ER stress and inflammatory programs, centered on *DDIT3* (CHOP), alongside upregulation of naive cell identity states in lymphocytes. Together, these observations suggest substantial plasticity in the aging immune landscape, even in individuals with established metabolic disease.

The coordinated downregulation of *DDIT3* (i.e., CHOP) across all immune lineages is the most robust finding of this study. CHOP, a downstream effector of the PERK arm of the unfolded protein response (UPR), is recognized as an amplifier of inflammatory and metabolic stress programs^62,63^. Prolonged ER stress can result in an irreversible cell state that triggers apoptosis through CHOP^64,65^ activation. While CHOP is not a primary initiator of inflammation, its sustained activation reinforces cytokine production, metabolic dysfunction, and immune stress^62^. Our data provide direct human evidence that *DDIT3* centered transcriptional programs are dynamic and can be attenuated through non-pharmacologic intervention, even in older adults with established metabolic disease. In CD14^+^ monocytes, *DDIT3* was co-downregulated with pro-inflammatory genes *OSM* and *CXCL8* within the same cells and were among the target genes in the inferred DDIT3 regulatory network, suggesting that DDIT3 may be an upstream regulator of attenuated inflammation in this context. Reduced *DDIT3* expression was also observed in CD8⁺ TEMRA and NK cells, consistent with the pan-lineage pattern. Prior work in tumor models has shown that deletion of CHOP in T cells enhances CD8⁺ effector function^66^, suggesting that reduced *DDIT3* may reflect improved adaptive immune cell function alongside reduced cellular stress. Consistent with this, CR has been shown to suppress the adipokine SPARC in humans, which drives pro-inflammatory IRF3/7-dependent interferon and cytokine programs in macrophages during aging and obesity, suggesting that adipose-derived metabolic signals may contribute to the attenuation of myeloid inflammatory programs observed here^67^.

Intervention was associated with upregulation of naive T and B cell identity genes across naive lymphocyte compartments, including *LEF1* and *TCF7*, which have previously been associated with a more stem-like phenotype^68,69^. Data from an independent healthy cohort^3^ showed that naiveness associated gene programs declined with age, suggesting that lifestyle intervention may partially reverse age-associated transcriptional programs in naive lymphocytes. RUNX1, whose regulon activity increased with intervention, has recently been associated with more youthful T cell signatures^70^, further supporting this interpretation. Loss of naive lymphocytes and accumulation of terminally differentiated cells are hallmarks of T cell aging that are exacerbated with obesity and metabolic disease^71^. Our findings suggest that transcriptional and epigenetic features of naive cell identity may be partially restored through CR and exercise, even in the presence of frailty. Prior work has shown that high levels of physical activity in older adults are associated with preserved naive T cell frequencies and reduced markers of immunosenescence, supporting the possibility that the exercise component of our intervention may have contributed to the transcriptional restoration of naive cell identity programs observed here^72^. How these changes in naive T cell transcriptional programs affect their functionality warrant further investigation. There was a notable upregulation of *IRF3* expression and binding site accessibility specifically in Tfh cells. *IRF3* plays central roles in antiviral defense and interferon responses in immune cells^73^. Chronic inflammation with aging and obesity can impair interferon signaling and antiviral immunity^13,15,74–76^. Prior work has linked IRF activity and interferon-stimulated gene signatures to improved vaccine outcomes^15,77–79^, raising the hypothesis that lifestyle intervention may enhance adaptive immune responses, a possibility that warrants further investigation.

We also observed downregulation of ’mTORC1 signaling’ and ’MAPK signaling’ gene sets across lymphocyte compartments. Mechanistically, reduced nutrient availability suppresses insulin–IGF-1–dependent phosphatidylinositol 3-kinase (PI3K) and mitogen-activated protein kinase (MAPK) signaling, leading to decreased activation of mammalian target of rapamycin (mTOR)^30^. Attenuation of mTOR signaling promotes autophagy and cellular stress resilience, which together contribute to suppression of chronic inflammation through reduced inflammasome activity and diminished pro-inflammatory cytokine production^30^. Restraint of mTORC1 is required for naive T cell quiescence, and elevated mTORC1 activity known to disrupt naive T cell identity and homeostasis^80,81^. Whether downregulation of mTOR signaling contributes to the observed shift toward naive cell identity restoration requires further investigation. Pharmacologic mTOR inhibition has been shown to improve influenza vaccine responses in older adults^82^, raising the possibility that lifestyle-mediated attenuation of mTOR activity may have similar benefits.

The increase in BCAA metabolism transcriptional programs was the top mitochondria-associated metabolic signal, consistent with prior associations between impaired BCAA catabolism and obesity and T2D^83^. A closely analogous signature has been reported in skeletal muscle and subcutaneous adipose tissue after combined endurance and resistance exercise, where enhanced BCAA catabolic gene expression mediated improvements in insulin sensitivity^84^, suggesting that our parallel signature in circulating immune cells reflects a systemic program of restored BCAA catabolic capacity. Pro-inflammatory cytokines and UPR activation have been shown to directly suppress BCAA catabolic gene expression in metabolic tissues^85^, linking the *DDIT3* and BCAA metabolism transcriptional modules as interconnected responses to lifestyle intervention. Furthermore, while exogenous BCAA supplementation has been associated with pro-inflammatory macrophage polarization^44,86^, BCAA oxidation through catabolic enzymes is required for alternative macrophage polarization and oxidative phosphorylation^87^, consistent with our observation that upregulation of BCAA catabolic transcripts co-occurs with reduced NF-κB and TNFα signaling. The negative correlation between BCAA metabolism scores and TNFα signaling gene expression, and the positive correlation with insulin sensitivity, further support a model in which intervention-induced metabolic reprogramming contributes to attenuation of chronic immune activation.

Despite these findings, several limitations warrant consideration. The cohort is modest in size (*n* = 9), which limits statistical power, though this is partially mitigated by the longitudinal paired design. While transcriptional and epigenetic changes suggest altered immune states, we did not directly assess functional immune outcomes such as vaccine immunogenicity or pathogen challenge, given the difficulty of conducting such studies in this population. Our analyses focused on circulating immune cells; whether similar remodeling occurs in tissue-resident immune compartments remains an open question. Finally, our analysis of BCAA metabolism is based on transcriptional pathway scoring and does not directly measure BCAA flux or enzyme activity. Future studies integrating controlled metabolic interventions with functional immune assays, targeted metabolomics, and longer-term follow-up will be essential to build upon our study.

Lifestyle intervention in older adults with obesity and T2D is associated with coordinated remodeling of immune cell states across innate and adaptive compartments. By attenuating ER stress and inflammatory programs and enhancing naive lymphocyte identity associated transcriptional states, these findings reveal substantial plasticity of the aging immune system even in late life and with established metabolic disease. This work supports lifestyle-based interventions as a potent non-pharmacologic strategy for immune remodeling in aging populations, and highlights ER stress, inflammation, and naive lymphocyte identity programs as key modifiable programs.

## METHODS

### Study Design

The Lifestyle Intervention for Senior Diabetics (LISD) trial (ClinicalTrials.gov NCT02348801) involving community-dwelling older adults (> 65 years) with obesity and Type 2 Diabetes^28^ was utilized for longitudinal transcriptional (scRNA-seq) and epigenetic (snATAC-seq) profiling of peripheral blood mononuclear cells (PBMCs). Participants underwent a 6-month lifestyle intervention consisting of caloric restriction (500–750 kcal/day deficit) combined with exercise training at Baylor College of Medicine and the Michael E. DeBakey VA Medical Center between April 2015 and December 2020. Following IRB approval (IRB protocol 24X-068-1) for this ancillary study involving additional blood collection, ten consecutive participants were enrolled for PBMC analysis. Samples from one participant was excluded from downstream analyses due to sample processing related issues. The final dataset comprised scRNA-seq data from 9 participants (n = 18 samples) and snATAC-seq from 5 participants (n = 10 samples). Further details on the lifestyle intervention and the cohort, including detailed inclusion and exclusion criteria, can be found in the parent study^28^.

### Glycemic Control and Insulin Sensitivity Assessment

HbA1c and Matsuda index values were assessed as previously described^28,88^. Briefly, HbA1c was measured using with Tosoh Automated Glycohemoglobin Analyzer (HLC-723G8; Tosoh Bioscience, San Francisco, CA) in the morning after hours of fasting^28^. Insulin sensitivity was estimated from the oral glucose tolerance test using the Matsuda index. Fasting and post-load plasma glucose and insulin concentrations were entered into the standard Matsuda formula to derive a whole-body insulin sensitivity index, with higher values reflecting greater insulin sensitivity^88^.

### Frailty Index

Frailty was assessed at baseline using a deficit-accumulation Frailty Index^28,34^. This index was constructed from a set of age-related symptoms, comorbidities, functional limitations, and abnormal clinical or laboratory findings, and was calculated as the proportion of deficits present out of the total number considered (range 0–1, with higher scores indicating greater frailty). Based on previous frailty categorization^35^, participants in this study were classified as pre-frail (FI > 0.1 and FI ≤ 0.21) or frail (FI > 0.21and ≤ 0.45).

### Physical Performance Test (PPT)

Physical function was measured with the Physical Performance Test (PPT)^28,89^. Participants completed a series of standardized tasks (e.g., walking, chair rises, lifting, stair climbing), each scored on an ordinal scale and summed to a total PPT score, where higher scores indicate better physical performance^28,89^.

### Reactive oxygen species byproduct measurement in serum

Oxidative stress was assessed by measuring serum 8-isoprostane (8-epi-PGF2α), a marker of lipid peroxidation^36^, using a competitive-ELISA kit (abx257156, Abbexa). Sample preparation and all test procedures were performed according to the manufacturer’s instructions. The concentration of 8-iso was expressed as picograms per milliliter.

### Sample preparation and processing for scRNA-seq

PBMCs were thawed quickly at 37oC and transferred into DMEM supplemented with 10% FBS. Cells were quickly spun down at 400 g, for 10 min. Cells were washed once with 1 x PBS supplemented with 0.04% BSA and finally re-suspended in 1 x PBS with 0.04% BSA. Viability was determined using trypan blue staining and measured on a Countess FLII. Briefly, 12,000 cells were loaded for capture onto the Chromium platform using the single cell 3’ gene expression reagent kit (v3 or v3.1) (10x Genomics). Following capture and lysis via Chromium Chip G, cDNA was synthesized and amplified (12 cycles) as per manufacturer’s protocol (10x Genomics, protocols CG000204; CG000315). Amplified cDNA and libraries were checked for quality on Agilent 4200 Tapestation, quantified by KAPA qPCR, and sequenced on an Illumina NovaSeq 6000 S4 flow cell (28-10-10-90 asymmetric configuration) targeting 50,000 raw read pairs per cell.

### Sample preparation and processing for snATAC-seq

For single nucleus ATAC sequencing (snATAC-seq) experiments, viable single cell suspensions from each sample were used to generate snATAC-seq data using the 10x Chromium platform according to the manufacturer’s protocols (10x Genomics, protocols CG000169; CG000168). Briefly, >100,000 cells from each sample were centrifuged and the supernatant was removed without disrupting the cell pellet. Lysis Buffer was added for 5 minutes on ice to generate isolated and permeabilized nuclei, and the lysis reaction was quenched by dilution with Wash Buffer. After centrifugation to collect the washed nuclei, diluted Nuclei Buffer was used to re-suspend nuclei at the desired nuclei concentration as determined using a Countess II FL Automated Cell Counter and combined with ATAC Buffer and ATAC Enzyme to form a Transposition Mix. Transposed nuclei were immediately combined with Barcoding Reagent, Reducing Agent B and Barcoding Enzyme and loaded onto a 10x Chromium Chip H for droplet generation followed by library construction. The barcoded sequencing libraries were subjected to bead clean-up and checked for quality on an Agilent 4200 TapeStation, quantified by KAPA qPCR, and pooled for sequencing on an Illumina NovaSeq 6000 S4 flow cell with recommended sequencing configuration (50-8-16-50) and trimmed for 2x50bp libraries.

### scRNA-seq data processing

Reads from scRNA-seq were aligned to 10x Genomics GRCh38 reference 2020-A and processed using Cell Ranger v7.1.0^90^. Ambient RNA was corrected using SoupX version 1.6.2^91^. Multiplets were detected and removed using Scrublet^92^ with default parameters: doublet rate=0.06, min counts=2, min cells=3, min gene variability PCTL=85, number of PCs=30. High quality cells were selected in each sample separately using the following criteria: 1) % mitochondria < 20%; 2) number of genes detected >= 200. Reads were then log normalized using *normalize_total* (target_sum=1e4) followed by *log1p* in SCANPY^93^. Data from each sample was then concatenated to form a single merged object for scRNA-seq.

### scRNA-seq clustering and annotation

The merged scRNA-seq object was first scaled: *scale(max_value=10)*. Highly variable genes were calculated using:*highly_variable_genes* in SCANPY^93^ using batch as the batch key. Initial clustering of scRNA-seq data was performed using principal component analysis (PCA): *PCA (svd_solver=’arpack’, n_comps=100)*, followed by batch correction using Harmony^94^ in SCANPY^93^: (*harmony_integrate*). Nearest neighbors were identified using *neighbors* in SCANPY^93^. Clusters were identified using Leiden (resolution = 1) and visualized using uniform manifold approximation and projection (UMAP). Red blood cell clusters (high *HBB* expression and low number of genes detected) were removed after identifying erythroblasts (high expression of *ALAS2*). Clusters assigned to major immune lineages were iteratively re-clustered, adjusting the number of principal components and Leiden clustering resolution parameters as necessary to identify known subsets within the respective lineage. Re-clustering was performed by recalculating nearest neighbors and applying UMAP for visualization. Further subclustering was performed as needed to clearly define known subsets. Subclusters exhibiting co-expression of gene expression markers from multiple immune cell lineages (i.e., multiplets) or elevated mitochondrial read fractions relative to neighboring subclusters were removed. Finally, annotations were assigned to clusters based on known marker gene expression.

### Memory CD4^+^ T Cell Annotation

Given the low expression of some memory CD4^+^ T cell subset marker genes, we annotated memory CD4^+^ T cells using Seurat (version 5.3.0)^95^ and Nebulosa (version 1.18.0)^96^. Read count matrices from memory CD4+ T cells were used as input into Seurat. Variable genes were identified using *FindVariableFeatures(selection.method = ’vst’, nfeatures = 2000)*. PCA was calculated using *RunPCA* using 50 principal components followed by *FindNeighbors* and *RunUMAP* functions to calculate nearest neighbors and visualize clusters. Harmony (version 1.2.3)^94^ was applied to correct for batch effects, followed by nearest neighbor calculation, Leiden clustering using *FindClusters*(*resolution=1, algorithm=’leiden’)*, and UMAP. Memory CD4^+^ T cell clusters were iteratively re-clustered as necessary to identify well defined memory CD4^+^ T cell annotations with known marker genes, using Nebulosa to amplify gene expression signatures. Final annotations and UMAP projections were saved and converted for further downstream analysis using SCANPY^93^ .

### snATAC-seq data processing

Reads from snATAC-seq were aligned to 10x Genomics GRCh38 reference 2020-A and processed using Cell Ranger ATAC v2.1.0^97^. High quality nuclei were selected based on the following criteria: % of reads within exclusion list regions < 5%, nucleosome signal < 4, % of fragments in peaks > 15%, number of peak region fragments > 3000, % of fragments at transcription start sites > 10%, and % mitochondrial read fragments < 10%. After filtering for high quality nuclei, AMULET (version 1.1)^98^ was applied to each sample independently to identify multiplets using FDR < 5%. Multiplets were used to identify multiplet rich clusters for exclusion during downstream clustering analysis.

Sample data was merged by first identifying a unified peak set across all samples. Individual sample peaks from Cell Ranger ATAC were merged based on 1 base pair (bp) overlap. Peaks were excluded using the following criteria: peak length < 20bp; > 10,000bp, chrY peak, or represented by less than 3 samples. Read count matrices were generated based on the unified peak set and concatenated into one matrix for all nuclei across samples. Read counts were normalized using a reimplementation of the term frequency inverse document frequency (TF-IDF) normalization method described in Signac^99^. AnnData objects were generated using SCANPY^93^ from this matrix to apply truncated singular value decomposition (SVD) using *pca(zero_center=False)* method in SCANPY^93^. In parallel, gene activity scores were quantified using a Python reimplementation of gene activity score calculations in Signac^99^. Briefly, transcription start sites and termination sites from UCSC hg38 refflat database^100^ were used as a reference to quantify reads within the gene body and 2000bp upstream from the transcription start site. To account for batch effects, batch correction was applied to nearest neighbors using BBKNN^101^.

### snATAC-seq clustering and annotation

Batch corrected objects were clustered using Leiden (resolution = 1). Clusters with high frequencies of multiplets detected using AMULET (version 1.1)^98^ were discarded from further analysis. Batch corrected nearest neighbors were recomputed and clusters were separated into T, NK, B, and myeloid subsets based on gene activity scores. Each subset was subclustered by recalculated batch corrected neighbors using BBKNN^101^ on nuclei within the subset. Multiplet rich clusters and remaining nuclei identified as multiplets by AMULET^98^ were removed. Peaks were called using MACS2 (version 2.2.7.1)^102^ (-f BAMPE, --keep-dup all, -nomodel) in each lineage using the aggregated read counts from all samples and nuclei within the lineage. Data processing steps for TF-IDF normalization and truncated SVD were repeated for each lineage to create lineage-specific AnnData objects in SCANPY^93^.

Corresponding lineages from scRNA-seq objects were extracted and both scRNA-seq and snATAC-seq objects were converted to Seurat objects. Label transfer was performed on each lineage using *FindTransferAnchors(reduction=’CCA’)* and *TransferData* functions in Signac (version 1.7.0)^99^ and and Seurat (version 4.1.1)^95^. Subclusters identified within each lineage were assigned corresponding labels from scRNA-seq based on the frequency of predicted annotations and gene activity scores of known marker genes.

### Differential gene expression

Pseudobulk count matrices were generated for each level of annotation: total PBMCs, major immune lineages, and lineage specific subsets, using ADPBulk (https://github.com/noamteyssier/adpbulk). Genes were filtered such that any gene with a detection fraction below 5% in more than 5 donors was excluded. Donor samples were excluded if the minimum number of cells for the corresponding immune cell subset was less than 50 for either corresponding sample pair. Aggregated raw gene expression count matrices were then normalized using the calcNormFactors function in edgeR (version 3.36.0)^103–105^. Differential expression analyses were performed using a paired design, with donor pair included as a blocking factor in the model matrix. Dispersion was estimated using estimateDisp and model fitting was performed using glmQLFit, with differential expression between post- and pre-intervention conditions tested using glmQLFTest. P-values were corrected for multiple comparisons using the Benjamini-Hochberg procedure, and genes with FDR < 5% and |log fold change| > 0.4 were considered differentially expressed.

For heatmap visualization, raw counts were normalized using calcNormFactors and log₂-CPM values computed with cpm(log=TRUE, prior.count=1) in edgeR. Donor-pair batch effects were removed using ComBat (sva R package, parametric prior)^106^, after which values were row-scaled (z-scored).

### Gene set enrichment

Gene set enrichment analysis (GSEA) was performed using the fgsea package^107^. Gene sets were obtained from MSigDB (version 7.4) and included Hallmark, KEGG, and pathway interaction database (PID) gene set collections^39,40,108,109^. Additional gene sets included exhausted versus memory CD8+ T cell signatures (GSE9650), a senescence gene set (SenMayo)^110^, and naive CD4+ T cell and naive B cell identity signatures derived from an independent cohort of healthy individuals (described below). For each cell type, genes were ranked using a score computed as -log(pvalue) × sign(log fold change). GSEA was then performed on this ranked gene list against all gene sets using the fgsea Multilevel function. Gene sets with FDR < 5% were considered significantly enriched.

### Derivation of Naive CD4+ T Cell and Naive B Cell Identity Signatures

Naive cell identity gene signatures were derived from scRNA-seq data of an independent cohort of 95 healthy individuals^3^. For the naive CD4^+^ T cell signature, cells annotated as naive CD4^+^ T cells (CD4_Naive and CD4_Naive_SOX4) were compared against all other CD4^+^ T cell subsets (central memory, Tfh, T_h_1, T_h_2, T_h_17, T_EMRA_, HLA-DR^+^, and ISG^hi^). For the naive B cell signature, naive B cells were compared against all other B cell subsets. Pseudobulk count matrices were generated per donor and group using ADPBulk. Genes were excluded if their detection fraction fell below 5% in more than 5 donors within the naive cell population. Since each donor contributes cells to both the naive and non-naive groups, donor was included as a blocking factor in the EdgeR model matrix to control for inter-donor variability. Genes with FDR < 5% and log fold change > 0.5 were retained as the naive identity signature for each lineage and used as gene sets in subsequent GSEA analyses. These gene sets are shared in Supplementary Table 5.

Naiveness scores were computed as the mean expression of signature genes detected in ≥5% of cells per cell type. For visualization of naiveness scores across age in the cohort of healthy individuals, expression was batch-corrected across 10x Genomics chemistry versions using sc.pp.combat in Scanpy prior to score computation.

### Mitochondria pathway prioritization scoring

Donor corrected transcripts per million (TPM) read counts were obtained from scRNA-seq objects for each respective subset. Raw read counts were pseudo-bulked by sample using ADPBulk and converted to TPM values using *RPKM* in edgeR (version 4.6.2)^103–105^ and scaled to 1,000,000 transcripts for each sample. TPM values were donor corrected with *removeBatchEffect* in limma^111^, using donor identifier as the batch variable. Mitochondrial pathway prioritization scores^41^ were then obtained using mitoPPS scripts available from https://github.com/annamonzel/mitotyping in conjunction with the human MitoCarta 3.0 gene lists^112^.

### Differentially accessible peaks

Reads were aggregated for each subset, and peaks were called using MACS2 (version 2.2.7.1)^102^ (-f BAMPE, --keep-dup all, -nomodel). A peak-by-sample read count matrix was then generated by quantifying fragments overlapping the called peaks. Donor samples were excluded if the minimum number of nuclei for the corresponding immune cell subset was less than 50 for either sample in the pair. Peaks were filtered based on within-sample detection, defined as the presence of at least one fragment in at least 10% of nuclei. Peaks were retained if they met this detection criterion in at least 33% of samples. The remaining peaks were further filtered using *filterByExpr* in edgeR(version 4.6.2)^103–105^, followed by normalization using *calcNormFactors*. Differentially accessible peaks were identified using a quasi-likelihood negative binomial model (*estimateDisp* and *glmQLFit*), with donor identity as a covariate (∼donor id + group). P-values were adjusted for multiple testing using the Benjamini-Hochberg procedure. Differentially accessible peaks are provided in Supplementary Table 8.

### Data integration with SCENIC+

Peaks were called using aggregated reads from all samples for each subset with MACS2 (version 2.2.7.1)^102^ (-f BAMPE, --keep-dup all, -nomodel). Peak and fragments files were used to generate cisTopic (version 2.0a0)^113^ objects using *create_cistopic_object_from_fragments*. The optimal number of topics for each subset was estimated using Mallet (version 2.0)^51^ with *run_cgs_models_mallet*, testing the following number of topics: 2, 5, 10, 15, 20, 25, 30, 35, 40, 45, 50. Optimal number of topics selected for each subset were as follows: Adaptive NK: 15, CD14^+^ monocyte: 25, CD16^+^ monocyte: 45, CD4^+^ T_EMRA_: 25, CD56^bright^ NK: 15, CD56^dim^ NK: 25, GZMK+ CD8+ T: 30, CD8^+^ T_EMRA_: 30, DC: 20, γδ T: 20, GZMK^+^ GZMH^+^ CD8^+^ T: 20, memory B: 25, memory CD4^+^ T: 40, naive B 20, naive CD4^+^ T: 25 naive CD8^+^ T: 15, Treg: 20, and pDC: 10. Region sets were defined using the top 3,000 regions per topic (ntop) and Otsu thresholding (otsu) as the default input to SCENIC+^51^. Region sets derived from differentially accessible peaks were additionally included when compatible with SCENIC+ execution; otherwise, analyses proceeded using the default region sets (ntop and Otsu). Corresponding scRNA-seq data was prepared with non-normalized and non-scaled read counts as required for SCENIC+. A cisTarget (version 1.1)^51^ database was generated for each subset using create_cistarget_motif_databases.py in SCENIC+^51^ with 1kb padded hg38 reference sequences at subset peaks and human motifs (V10: 2022 SCENIC+ clustered motif collection) downloaded from: https://resources.aertslab.org/cistarget/motif_collections/v10nr_clust_public/v10nr_clust_public.zip. Finally, SCENIC+ (version 1.0a2)^51^ was run using default parameters (min_target_genes: 10, ctx_auc_threshold: 0.005, ctx_nes_threshold: 3.0, ctx_rank_threshold: 0.05, dem_log2fc_thr: 1.0), with *is_multiome=False* and 10 cells per meta-cell.

### Transcription factor motif binding site accessibility

Reads were aggregated for each subset, and peaks were called using MACS2 (version 2.2.7.1)^102^ (-f BAMPE, --keep-dup all, -nomodel). Read count matrices were generated from regions centered on peak summits and extended ±250 bp. Transcription factor (TF) binding site accessibility deviation scores were computed using *computeDeviations* in chromVAR version 1.16.0^58^ with motifs from the JASPAR 2024 CORE database^114^. TFs were prioritized based on consistency of paired shifts (Wilcoxon signed-rank test) and differences in distributions (Mann–Whitney U test) of median deviation scores between conditions, using nominal p-value thresholds (0.0625 and 0.05, respectively) as heuristic filters for prioritization and ranking.

### Co-expression analysis

For each gene pair, the number of cells expressing each gene was quantified based on log-normalized expression values > 0. Cells co-expressing both genes were defined as double-positive cells. Co-expression frequency was calculated as the proportion of double-positive cells within each subset.

### Statistical tests

Paired comparisons between donor time points were performed using the Wilcoxon signed-rank test (*wilcoxon* in the SciPy Python package), unless otherwise noted. Correlations between variables were calculated using Spearman’s rank correlation coefficient (*spearmanr* in SciPy). Mann-Whitney U test (mannwhitneyu in SciPy) was used to assess differences in regulon activity scores between pooled meta-cell groups. Error bars for double-positive cell frequency bar plots represent standard deviation (*errorbar=“sd”* in Seaborn). Benjamini-Hochberg correction was applied for multiple hypothesis testing where applicable (*p.adjust* in R and *fdrcorrection* in statsmodels.stats.multitest in Python), except for analyses where adjusted p-values were provided by the method itself (e.g., GSEA).

## COD AVAILABILITY

Code is available on github: https://github.com/UcarLab/Lifestyle

## AUTHOR CONTRIBUTIONS

L.F., J.B., D.T.V., G.A.K., D.U designed the study and raised the funds. D.T.V., G.A.K.,

D.U. co-supervised the study. R.Y. and A.T. led the data analyses. D.N-B., A.P.W., M.L.S., and S.P. helped with data analyses and data interpretation. R.M. generated genomics data. R.Y., A.T., and D.U. wrote the manuscript. All authors revised the manuscript and helped with data interpretation.

## ACKNOWLEDGEMENTS

We thank JAX Genomic Technologies and Single Cell cores for their help with generating the sequencing data. We thank Carmen Robinett for feedback and help with scientific writing. We thank Olivia Bart for help with dbGAP data upload. We thank members of the Ucar lab for critical feedback during the progress of the study. We thank the participants and LISD trial and the researchers who contributed to the prior work in the parent study. ChatGPT (OpenAI; GPT-5–series models) and Claude (Anthropic; Opus 4.6–4.7) were used for refinement, improving clarity of language, and early drafts for this manuscript.

---

**Extended Data Fig. 1.**
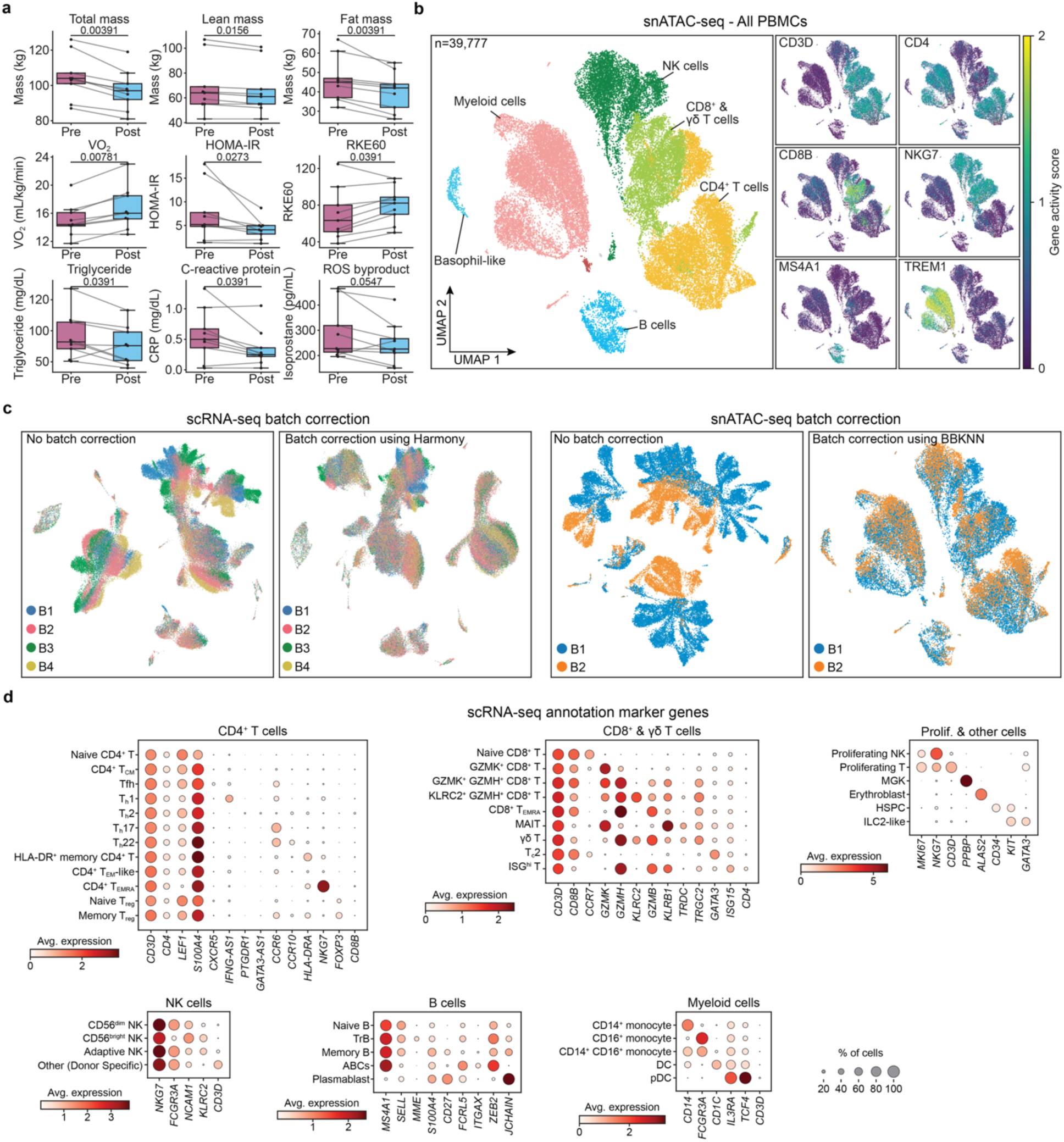
Clinical measurements, snATAC-seq overview, batch correction, and scRNA-seq annotation. **a**, Paired comparisons of additional clinical measures before and after intervention: total mass, lean mass, fat mass, VO₂, HOMA-IR, RKE60, triglycerides, C-reactive protein, and serum isoprostane levels. P values were calculated using the Wilcoxon signed-rank test. **b**, UMAP of 39,777 nuclei from snATAC-seq colored by annotated cell population (left) and feature plots of canonical marker gene activity scores (right). **c**, UMAPs showing scRNA-seq data before and after batch correction using Harmony (left) and snATAC-seq data before and after batch correction using BBKNN (right), colored by sequencing batch. **d**, Dot plots of canonical marker gene expression used for annotation of CD4⁺ T cell, CD8⁺ and γδ T cell, NK cell, B cell, myeloid cell, proliferating and other cell subsets in scRNA-seq data. Dot size reflects the percentage of cells expressing each gene; dot color reflects average expression.

**Extended Data Fig. 2.**
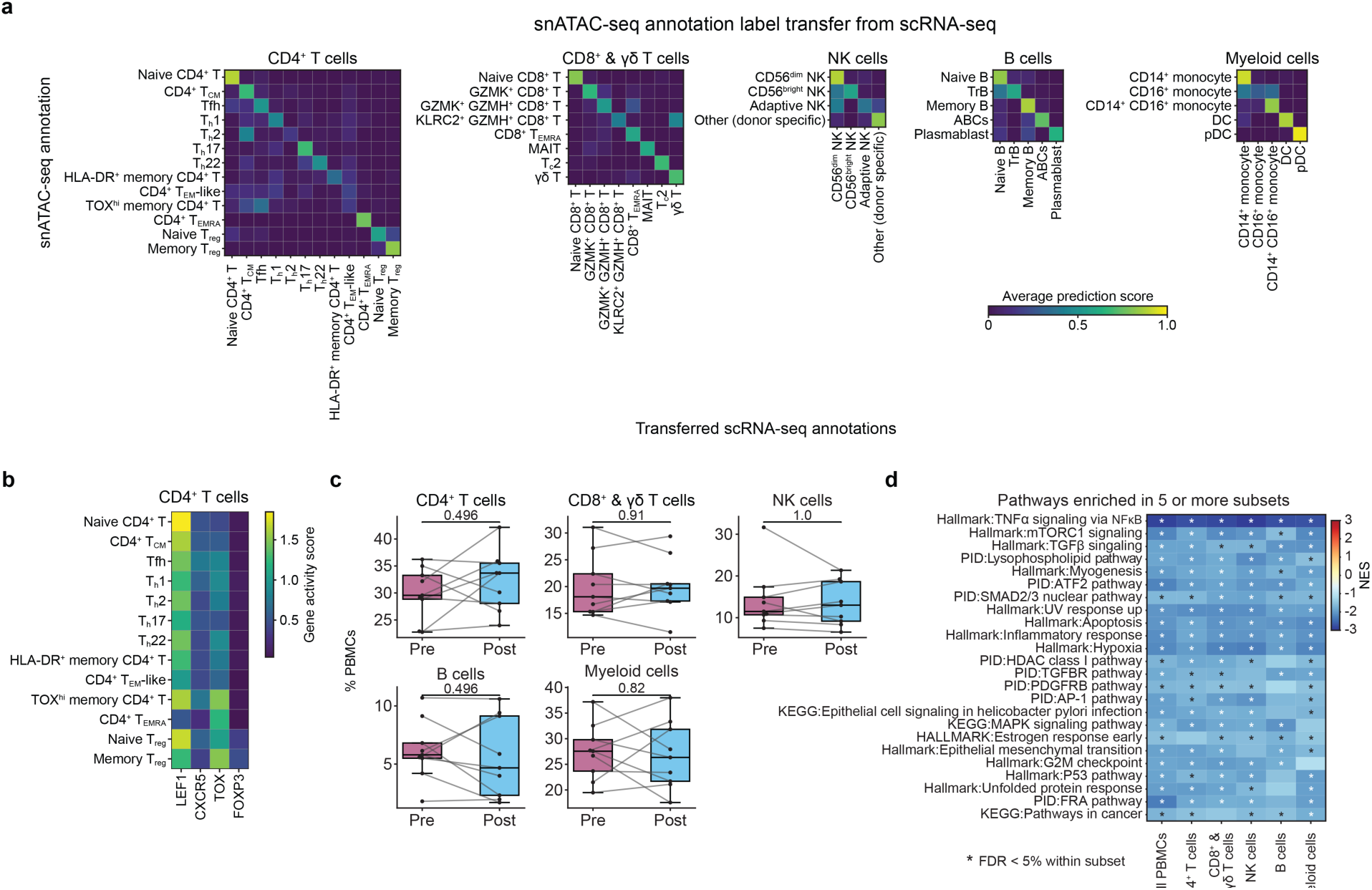
snATAC-seq label transfer, cell type frequencies, and pathway enrichments across PBMC lineages. **a**, Heatmaps of average label transfer prediction scores from scRNA-seq to snATAC-seq annotations across CD4⁺ T cell, CD8⁺ and γδ T cell, NK cell, B cell, and myeloid cell lineages. **b**, Heatmap of gene activity scores for select marker genes (LEF1, CXCR5, TOX, FOXP3) across CD4⁺ T cell subsets in snATAC-seq data. **c**, Paired comparisons of major PBMC lineage frequencies as a percentage of total PBMCs before and after intervention. P values were calculated using the Wilcoxon signed-rank test. **d**, Heatmap of normalized enrichment scores (NES) for gene sets enriched in five or more PBMC subsets following intervention, across total PBMCs and major lineages. Asterisks indicate FDR < 5% within the corresponding subset.

**Extended Data Fig. 3.**
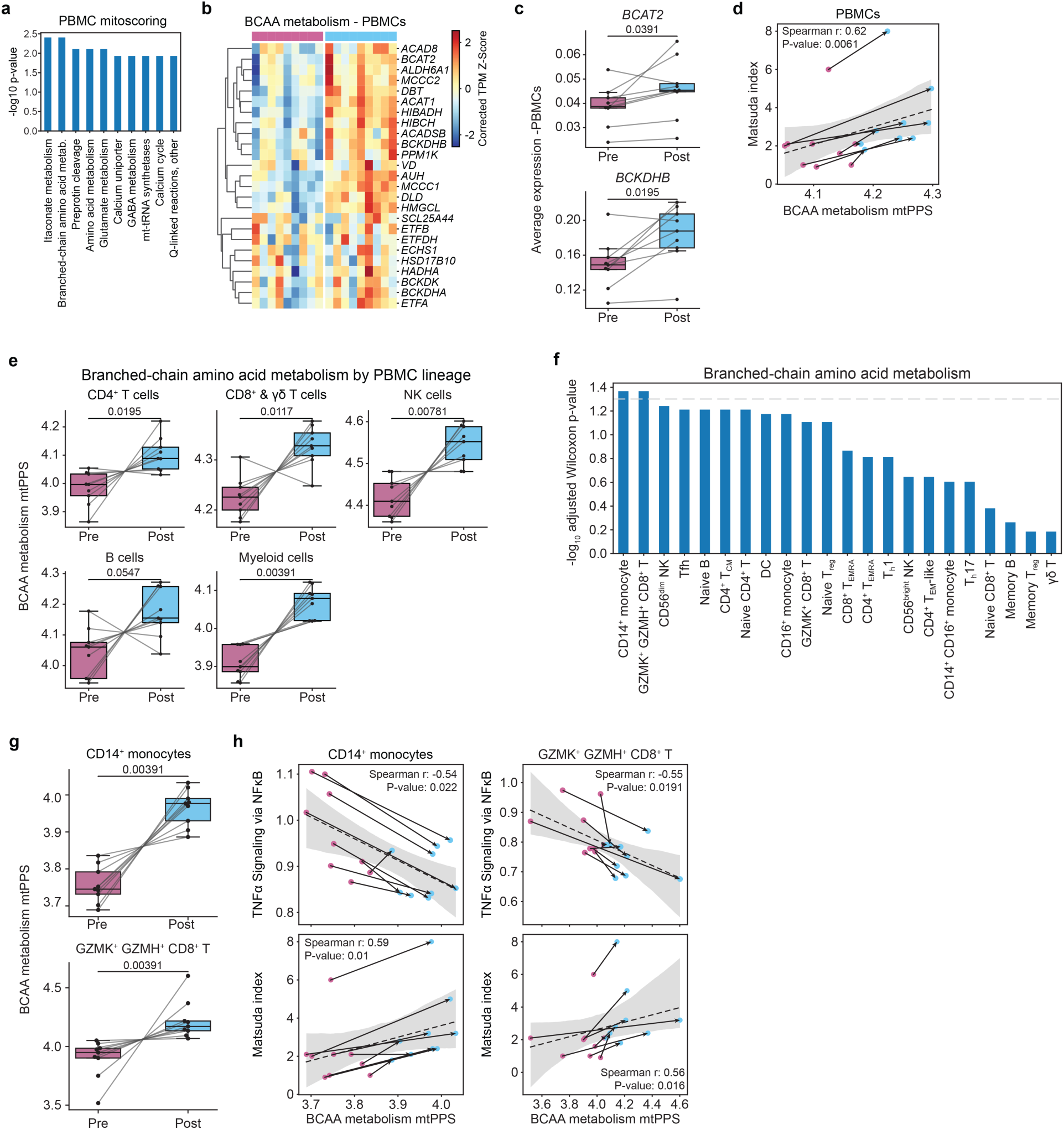
Branched-chain amino acid metabolism is upregulated following intervention across PBMC lineages. **a**, Bar plot of the top ten mitochondrial pathways for PBMCs, ranked by −log10 p-value comparing pre versus post mitochondrial pathway prioritization scores (mtPPS) using the the Wilcoxon signed-rank test. **b**, Heatmap of corrected TPM z-scores for BCAA metabolism genes across donors before and after intervention in total PBMCs. **c**, Average expression of *BCAT2* and *BCKDHB* in total PBMCs before and after intervention. **d**, Correlation between BCAA metabolism mtPPS and Matsuda index across donor timepoints in total PBMCs. **e,** BCAA metabolism mtPPS before and after intervention across major PBMC lineages. **f**, Bar plot of −log10 adjusted p-values for BCAA metabolism mtPPS changes across PBMC subsets. Dashed line indicates the adjusted p-value significance threshold. **g**, BCAA metabolism mtPPS before and after intervention in CD14⁺ monocytes and GZMK⁺ GZMH⁺ CD8⁺ T cells. **h**, Correlations between BCAA metabolism mtPPS and ’TNFα signaling via NF-κB’ leading-edge gene expression score (top) and Matsuda index (bottom) in CD14⁺ monocytes (left) and GZMK⁺ GZMH⁺ CD8⁺ T cells (right) across donor timepoints (pink, Pre; blue, Post). Correlations and p values shown in d and h were calculated using Spearman’s correlation coefficient. P values in c, e, and g were calculated using the Wilcoxon signed-rank test.

**Extended Data Fig. 4.**
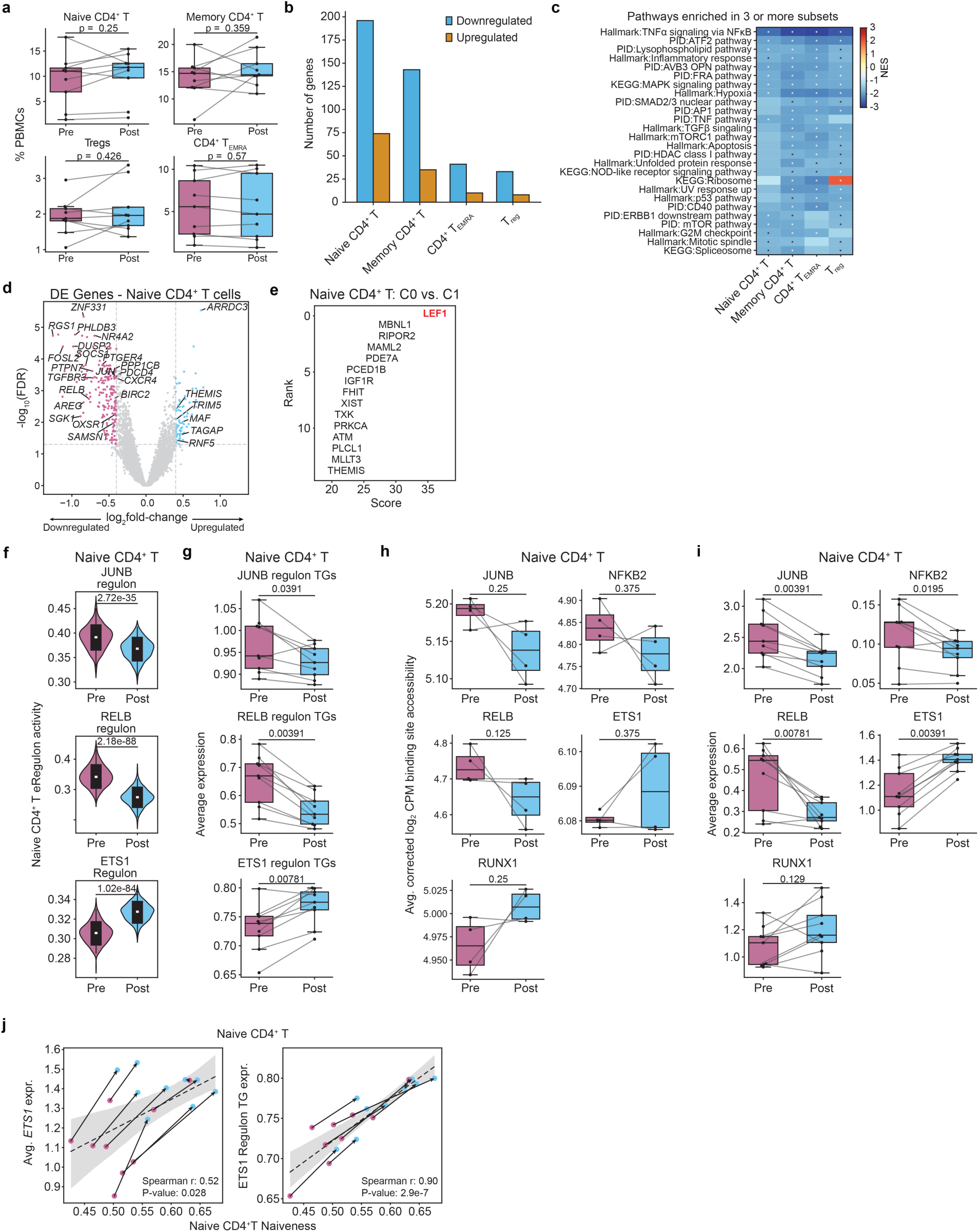
Naive CD4⁺ T cells have increased cell identity gene expression and increased RUNX1 activity in inferred regulatory networks. **a**, Frequencies of naive CD4⁺ T, memory CD4⁺ T, Treg, and CD4⁺ TEMRA cells as a percentage of total PBMCs before and after intervention. **b**, Number of upregulated and downregulated differentially expressed genes (FDR < 5%) across CD4⁺ T cell subsets. **c**, Heatmap of NES values for gene sets enriched in three or more CD4⁺ T cell subsets following intervention. Asterisks indicate FDR < 5% within the corresponding subset. **d**, Volcano plot of differentially expressed genes in naive CD4⁺ T cells (Post versus Pre). **e**, Ranked marker genes distinguishing naive CD4⁺ T cell Cluster 0 from Cluster 1, with *LEF1* highlighted as the top-ranked gene. **f**, Regulon activity for JUNB, RELB, and ETS1 regulons in naive CD4⁺ T cells before and after intervention. P values were calculated using the Mann–Whitney U test. **g**, Average expression of JUNB, RELB, and ETS1 regulon target genes in naive CD4⁺ T cells before and after intervention. **h**, Average corrected log2 CPM accessibility for inferred JUNB, NFKB2, RELB, ETS1, and RUNX1 binding sites in naive CD4⁺ T cells before and after intervention. **i**, Average expression of *JUNB*, *NFKB2*, *RELB*, *ETS1*, and *RUNX1* in naive CD4⁺ T cells before and after intervention. **j**, Correlations between naive CD4⁺ T cell naiveness scores and ETS1 expression (left) and ETS1 regulon target gene expression (right) across donor timepoints (pink, Pre; blue, Post). Correlations and p values were calculated using Spearman’s correlation coefficient. P values in a, g, h, and i were calculated using the Wilcoxon signed-rank test.

**Extended Data Fig. 5.**
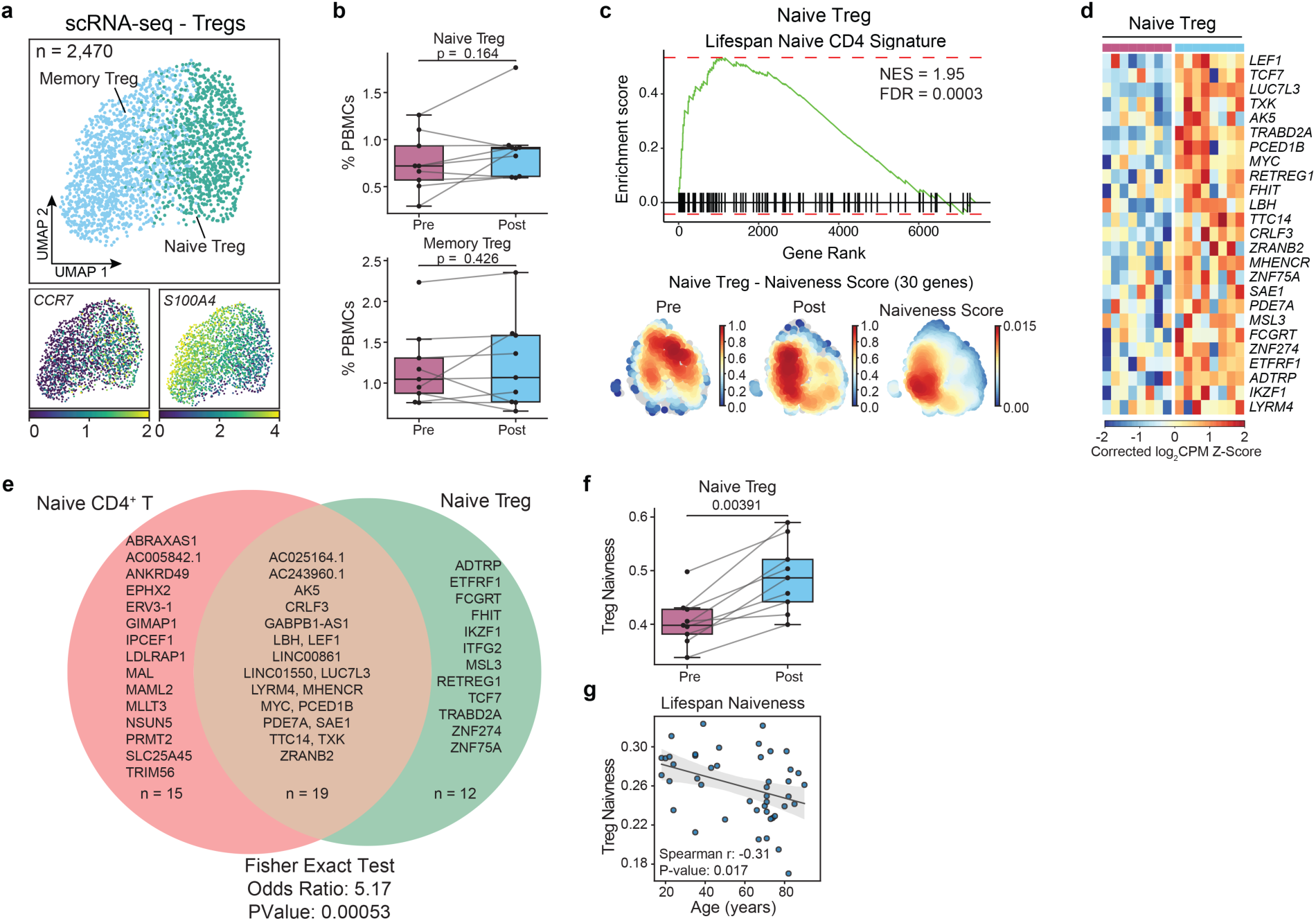
Naive Treg cells exhibit upregulation of naive cell identity genes following intervention. **a**, UMAP of 2,470 Treg cells colored by annotated subset (top) and feature plots of canonical marker genes (bottom). **b**, Frequencies of naive Treg and memory Treg cells as a percentage of total PBMCs before and after intervention. **c**, GSEA plot for the naive CD4⁺ T cell identity gene set in naive Treg cells (top) and density plots of naiveness scores in naive Treg cells before and after intervention (bottom). Normalized enrichment score (NES) and FDR from GSEA is shown. **d**, Heatmap of corrected log2 CPM z-scores for naiveness leading-edge genes across donors before and after intervention in naive Treg cells. **e**, Venn diagram showing overlap between naive CD4⁺ T cell and naive Treg naiveness leading-edge genes. Significance of overlap assessed by Fisher’s exact test; odds ratio and p value shown. **f**, Naiveness scores in naive Treg cells before and after intervention. **g**, Correlation between naive Treg naiveness scores and age in healthy adults (>18 years, n=45). Correlations and p values were calculated using Spearman’s correlation coefficient. P values in b and f were calculated using the Wilcoxon signed-rank test.

**Extended Data Fig. 6.**
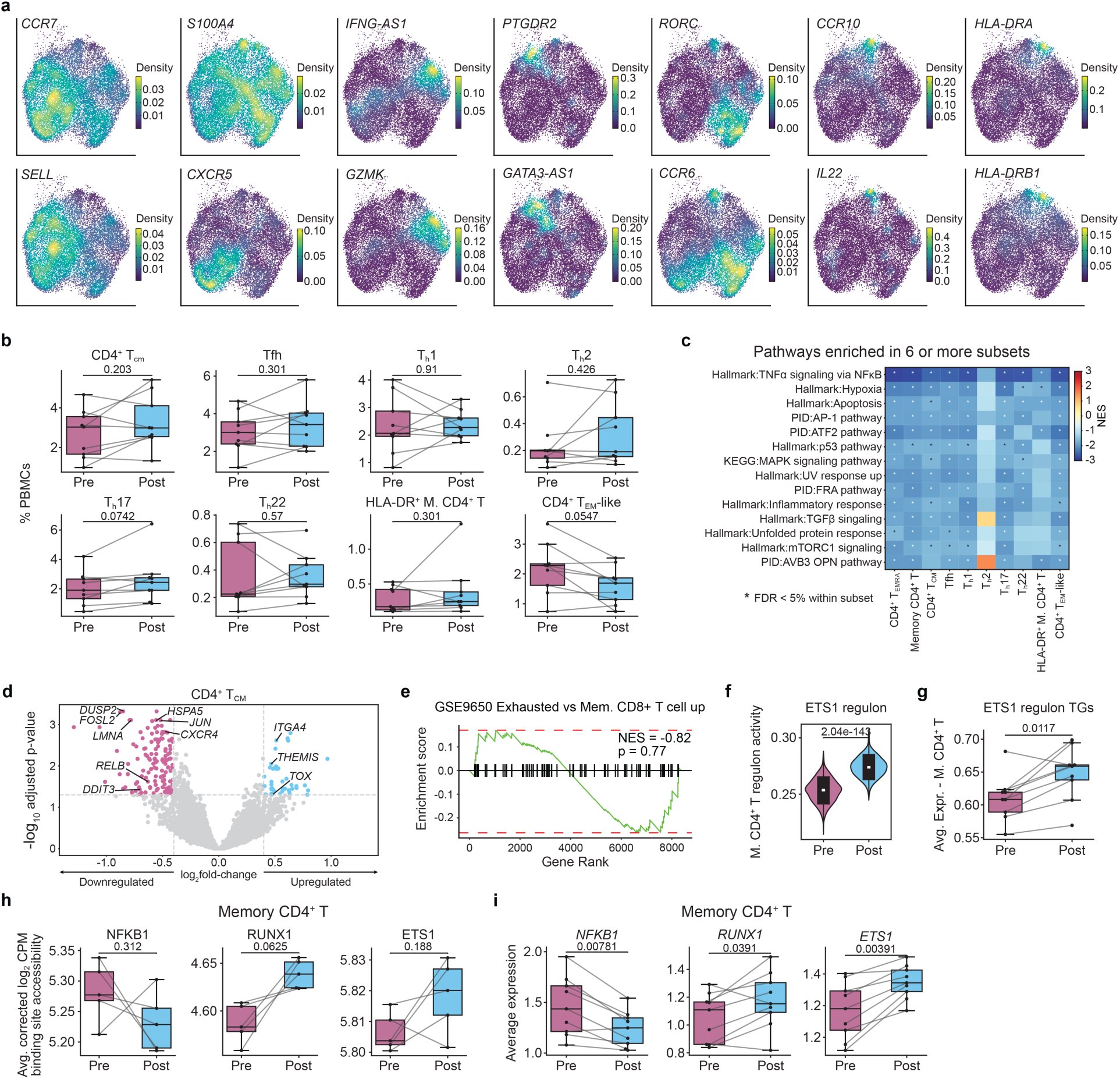
Memory CD4⁺ T cells have increased RUNX1 activity in inferred regulatory networks. **a**, Nebulosa density plots of canonical marker gene expression used for annotation of memory CD4⁺ T cell subsets. **b**, Frequencies of CD4⁺ TCM, Tfh, Th1, Th2, Th17, Th22, HLA-DR⁺ memory CD4⁺ T, and CD4⁺ TEM-like cells as a percentage of total PBMCs before and after intervention. **c**, Heatmap of NES values for gene sets enriched in six or more memory CD4⁺ T cell subsets following intervention. Asterisks indicate FDR < 5% within the corresponding subset. **d**, Volcano plot of differentially expressed genes in CD4⁺ TCM cells (Post versus Pre). **e**, GSEA plot for an exhausted versus memory CD8⁺ T cell upregulated gene set (GSE9650) in CD4⁺ TCM cells. Normalized enrichment score (NES) and p value shown. **f**, ETS1 regulon activity in memory CD4⁺ T cells before and after intervention. P value was calculated using the Mann–Whitney U test. **g**, Average expression of ETS1 regulon target genes in memory CD4⁺ T cells before and after intervention. **h**, Average corrected log_2_ CPM binding site accessibility for NFKB1, RUNX1, and ETS1 in memory CD4⁺ T cells before and after intervention. **i**, Average expression of *NFKB1*, *RUNX1*, and *ETS1* in memory CD4⁺ T cells before and after intervention. P values in b, g, h, and i were calculated using the Wilcoxon signed-rank test.

**Extended Data Fig. 7.**
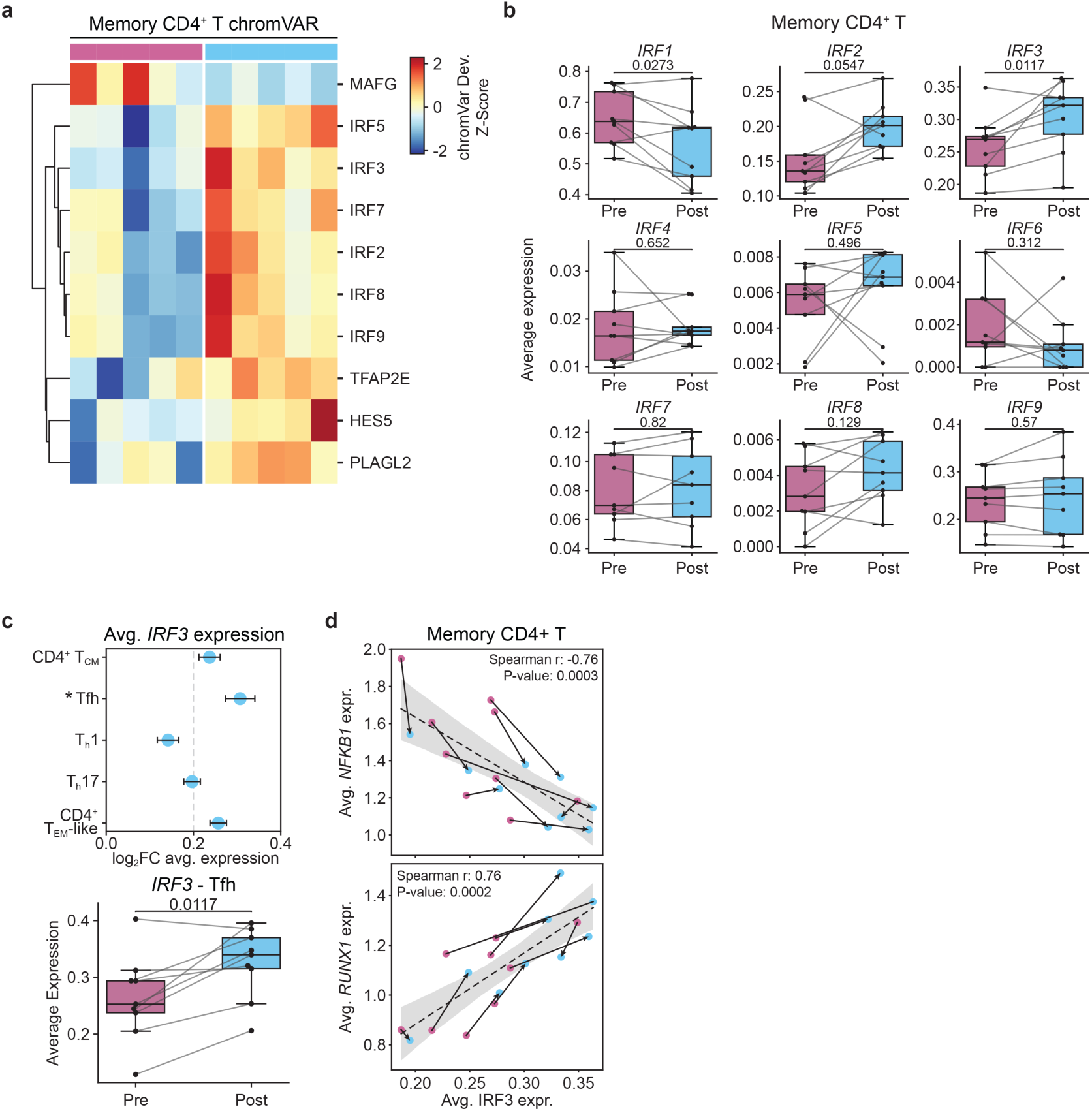
***IRF3* expression is increased in memory CD4⁺ T cells. a**, Heatmap of chromVAR deviation z-scores for top transcription factors in memory CD4⁺ T cells across donors before and after intervention. **b**, Average expression of IRF family members (*IRF1*-*IRF9*) in memory CD4⁺ T cells before and after intervention. **c**, Log_2_ fold change in average *IRF3* expression across memory CD4⁺ T cell subsets with asterisk indicating FDR < 5% (top) and paired comparison of *IRF3* expression specifically in Tfh cells before and after intervention (bottom). **d**, Correlations between average *IRF3* expression and average *NFKB1* expression (top) and average *RUNX1* expression (bottom) in memory CD4⁺ T cells across donor timepoints (pink, Pre; blue, Post). Correlations and p values were calculated using Spearman’s correlation coefficient. P values in b and c were calculated using the Wilcoxon signed-rank test.

**Extended Data Fig. 8.**
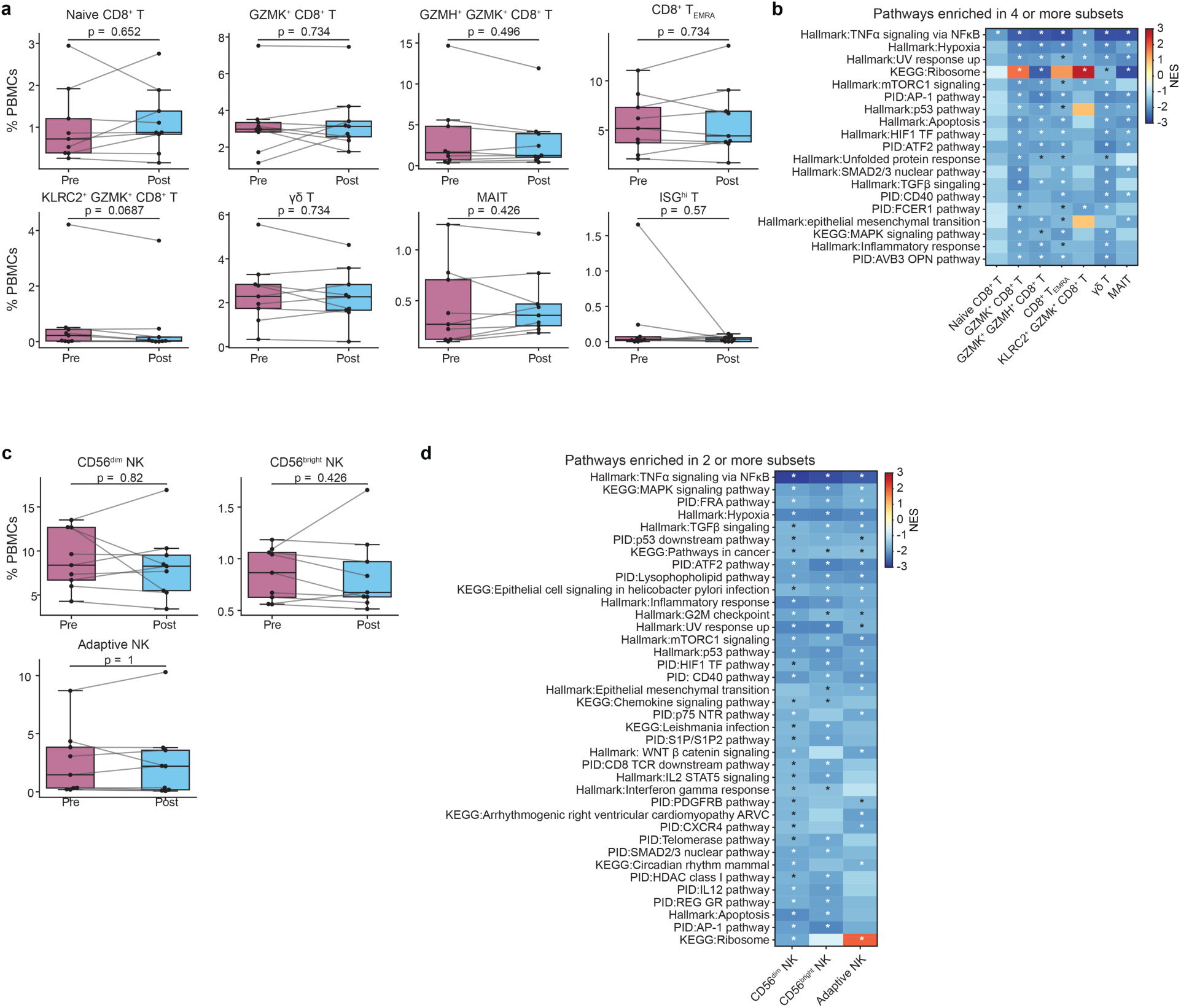
Cell type frequencies and pathway enrichments in CD8⁺ T and NK cell subsets. **a**, Frequencies of naive CD8⁺ T, GZMK⁺ CD8⁺ T, GZMK⁺ GZMH⁺ CD8⁺ T, CD8⁺ TEMRA, KLRC2⁺ GZMK⁺ CD8⁺ T, γδ T, MAIT, and ISGhi T cells as a percentage of total PBMCs before and after intervention. **b**, Heatmap of normalized enrichment scores (NES) for gene sets enriched in four or more CD8⁺ T cell subsets following intervention. Asterisks indicate FDR < 5% within the corresponding subset. **c**, Frequencies of CD56^dim^ NK, CD56^bright^ NK, and adaptive NK cells as a percentage of total PBMCs before and after intervention. **d**, Heatmap of NES values for gene sets enriched in two or more NK cell subsets following intervention. Asterisks indicate FDR < 5% within the corresponding subset. P values in a and c were calculated using the Wilcoxon signed-rank test.

**Extended Data Fig. 9.**
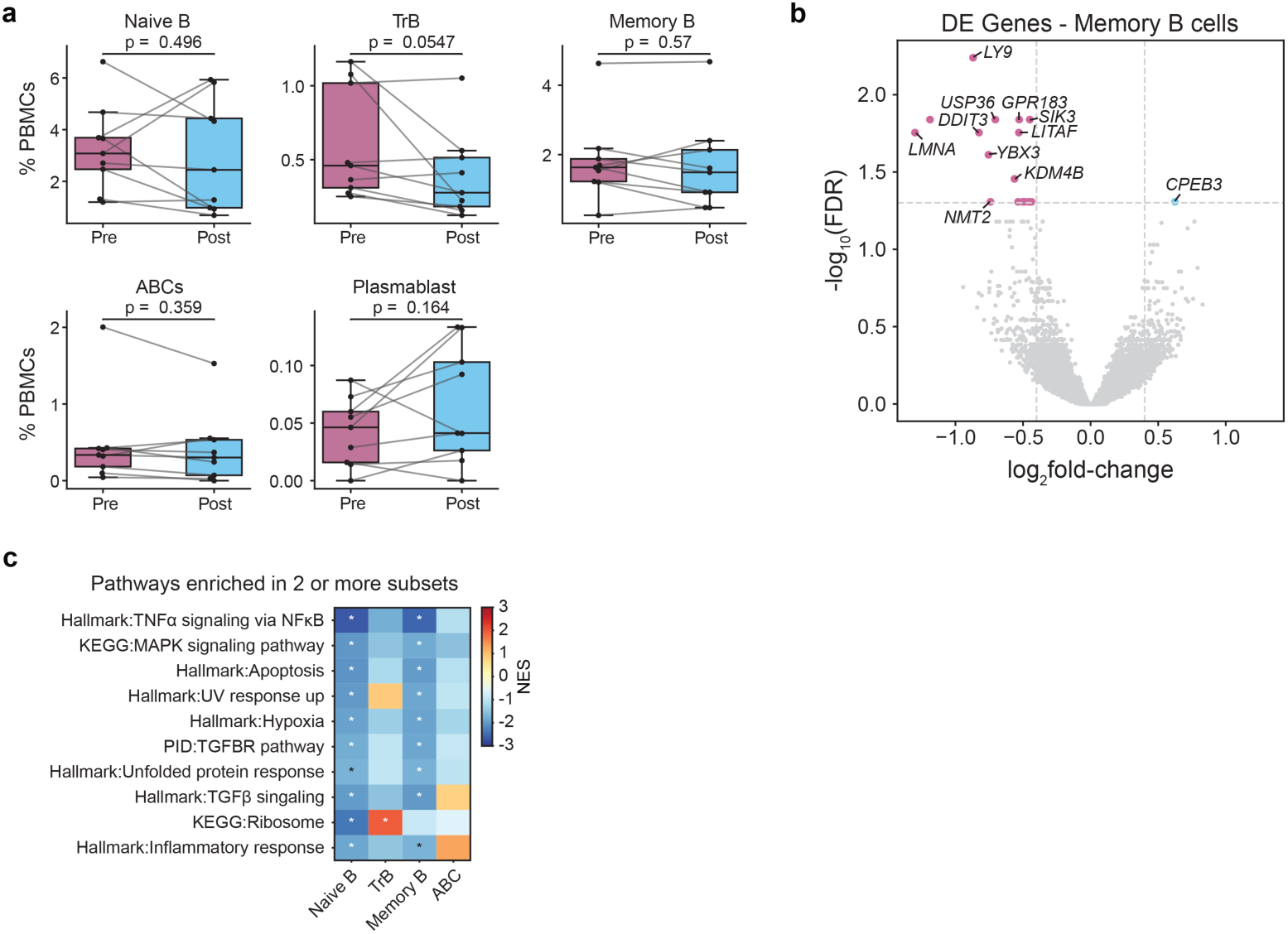
Cell type frequencies, differentially expressed genes, and pathway enrichments in B cell subsets. **a**, Frequencies of naive B, transitional B (TrB), memory B, ABC, and plasmablast cells as a percentage of total PBMCs before and after intervention. P values were calculated using the Wilcoxon signed-rank test. **b**, Volcano plot of differentially expressed genes in memory B cells (Post versus Pre). **c**, Heatmap of normalized enrichment scores (NES) for gene sets enriched in two or more B cell subsets following intervention. Asterisks indicate FDR < 5% within the corresponding subset.

**Extended Data Fig. 10.**
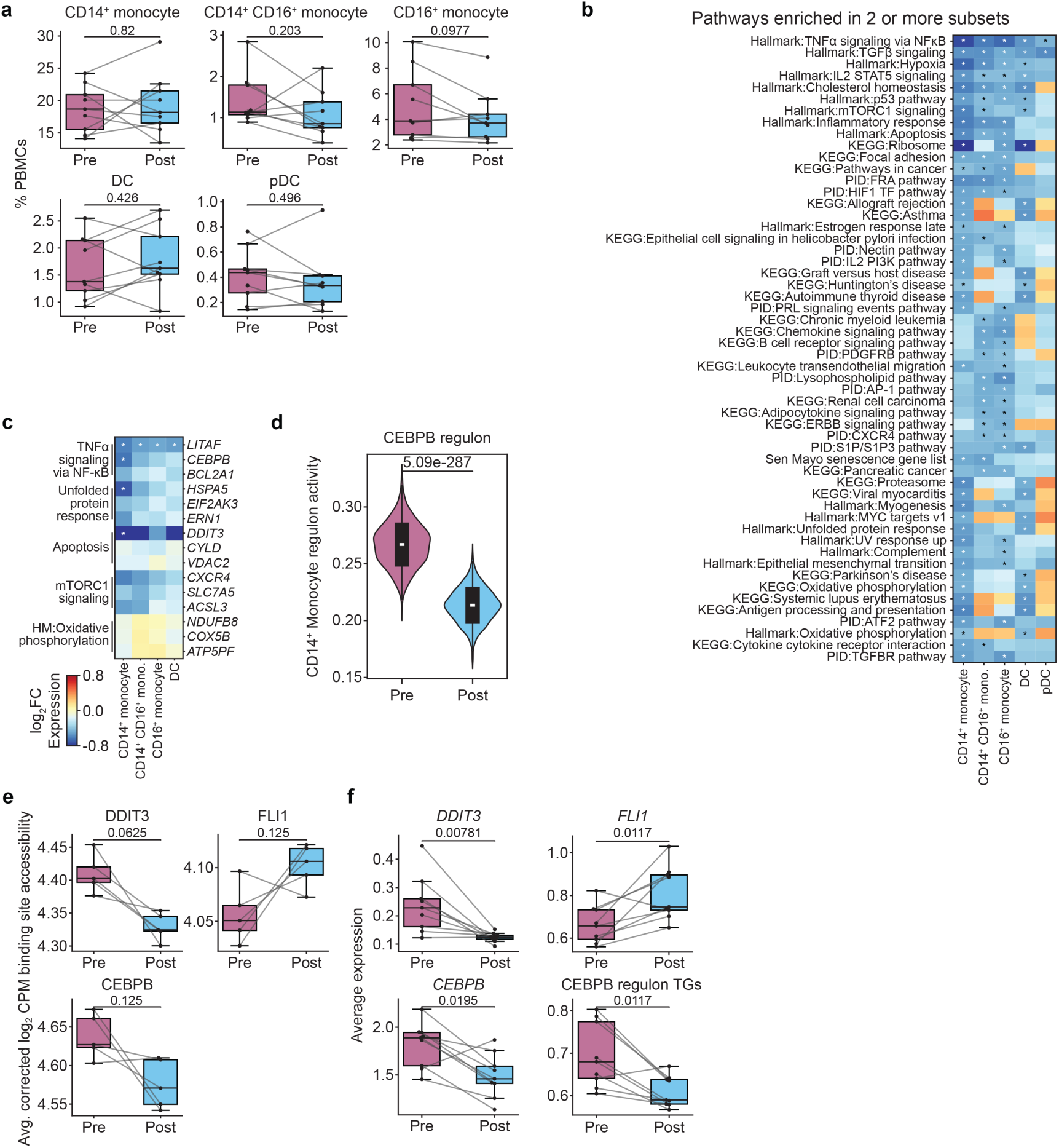
CD14⁺ monocytes have decreased DDIT3 and CEBPB activity in inferred regulatory networks. **a**, Frequencies of CD14⁺ monocyte, CD14⁺ CD16⁺ monocyte, CD16⁺ monocyte, DC, and pDC cells as a percentage of total PBMCs before and after intervention. **b**, Heatmap of normalized enrichment scores (NES) for gene sets enriched in two or more myeloid cell subsets following intervention. Asterisks indicate FDR < 5% within the corresponding subset. **c**, Heatmap of log_2_ fold changes (Post versus Pre) for selected differentially expressed genes across CD14⁺ monocyte, CD14⁺ CD16⁺ monocyte, and CD16⁺ monocyte subsets. Asterisks indicate FDR < 5%. **d**, CEBPB regulon activity in CD14⁺ monocytes before and after intervention. P was calculated using the Mann–Whitney U test. **e**, Average corrected log_2_ CPM binding site accessibility for *DDIT3*, *FLI1*, and *CEBPB* in CD14⁺ monocytes before and after intervention. **f**, Average expression of *DDIT3*, *FLI1*, and *CEBPB*, and average expression of CEBPB regulon target genes in CD14⁺ monocytes before and after intervention. P values in a, e, and f were calculated using the Wilcoxon signed-rank test.

